# ReCardioids: a robust human heart organoid platform to model doxorubicin- and radiotherapy-induced cardiotoxicity

**DOI:** 10.64898/2026.05.29.728705

**Authors:** Emil Rehnberg, Bjorn Baselet, Emre Etlioglu, Zoë Janssen, Ben Cools, Cynthia Van Rompay, Randy Vermeesen, Lorenzo Moroni, Sarah Baatout, Kevin Tabury

**Affiliations:** Department of Physics and Astronomy, Faculty of Sciences, Ghent University, Ghent, Belgium; Nuclear Medical Applications, Radiobiology unit, SCK CEN, 2400 Mol, Belgium; Department of Pharmaceutical and Pharmacological Sciences, Vrije Universiteit Brussel, Brussels, Belgium; Department of Biosystems, Mechatronics, Biostatistics and Sensors (MeBioS), Biomimetics group, KU Leuven, 3001 Leuven, Belgium; Department of Complex Tissue Regeneration, MERLN Institute for Technology-Inspired Regenerative Medicine, Maastricht, University, Maastricht, Netherlands

## Abstract

As global cancer incidence rises, so does the population of cancer survivors, making long-term and post-treatment quality of life a central clinical concern. Despite modern improvements in treatment, cardiotoxic side effects caused by anthracycline agents and thoracic radiotherapy remain a major clinical challenge. Human iPSC-derived heart organoids offer a promising alternative to animal and 2D *in vitro* models, but their utility is limited by inter-organoid variability. To address this, we developed reCardioids, a robust heart organoid model generated by dissociation and reaggregation of self-assembling cardioids. Through single-cell transcriptomics, we demonstrated that reCardiods maintain cellular diversity while also exhibiting a more mature cardiomyocyte phenotype compared to non-dissociated cardioids. To validate their utility as a preclinical *in vitro* model, we evaluated their response after exposure to doxorubicin and clinically relevant doses of γ-radiation. reCardioids successfully modeled doxorubicin-induced cytotoxicity, metabolic decline and altered contractile dynamics. Bulk RNA sequencing following radiation exposure revealed a temporal trajectory of injury, progressing from acute DNA damage through vascular stunting, metabolic dysfunction and eventually compensatory pathological hypertrophic remodeling.

## Introduction

While global cancer incidence continues to rise, advancements in early detection and targeted therapy have significantly improved survival, ^[1,2]^ thereby creating a growing population of cancer survivors. As a result, long-term and post-treatment quality of life, including cardiovascular health, has emerged as a central clinical concern.

A major complication of several standard cancer therapies is cardiotoxic side effects ^[3,4]^. Both chemo- and radiotherapeutic treatments can cause substantial cardiac toxicity that clinically resembles an accelerated aging phenotype ^[5–7]^. Anthracyclines such as doxorubicin are well known to cause dose-dependent cardiac dysfunction, ^[8,9]^ while thoracic radiotherapy, even with modern 3D conformal radiotherapy still subjects the heart to a substantial dose ^[10–12]^ that drives inflammation, progressive fibrosis and compensatory hypertrophy ^[13,14]^. Consequently, young cancer patients frequently develop cardiovascular diseases far earlier than the general population ^[15,16]^. Despite this clinical prevalence, few countermeasures, beyond dose limitation, are currently available ^[3,17]^.

Advances in understanding and mitigating these toxicities are hindered by the limitations of current preclinical models. Physiological responses to radiation or chemotherapeutics are heavily influenced by the complex multicellular interactions between different cell types, in particular cardiomyocytes, endothelial cells, pericytes and fibroblasts, ^[18–20]^ which are poorly captured by simple 2D culture systems. In addition, while *in vivo* models provide crucial systemic context, they often fail to predict individual human responses accurately ^[21,22]^. For example, mice tolerate much higher doses of doxorubicin than human patients and thus do not predict the toxicological effects of doxorubicin well ^[21,23,24]^, illustrating the concern regarding the use of animal models.

Human induced pluripotent stem cell (iPSC)-derived heart organoids offer a promising complementary platform, combining human-specific genotype and phenotype, with multicellular diversity and tissue-like microenvironment ^[25–30]^. While their use is expanding when modeling complex conditions and performing drug screening ^[31–33]^, their application in modeling radiation-induced heart disease remains scarce ^[21]^. Although a few recent studies have utilized advanced 3D engineered heart tissues (EHTs) to model radiation-induced dysfunction, these platforms typically rely on combining individually pre-differentiated cell types within an exogenous synthetic matrix ^[34–36]^. In contrast, developmental heart organoids undergo innate co-development and self-assembly from a multi-lineage progenitor state, establishing native microenvironments and highly integrated intercellular networks ^[25–30]^ that are essential for capturing the multicellular crosstalk triggered by ionizing radiation. While the very first proof-of-concept studies applying true heart organoids to radiobiology have recently emerged ^[37]^, the broader landscape of *in vitro* models still typically lacks this complex heterogenous architecture ^[34,35,38]^. Furthermore, existing *in vitro* radiation studies often rely on acute doses that far exceed the fractionated doses typically seen during clinical radiotherapy ^[37,39–41]^ and iPSC-derived organoids frequently suffer from high inter-organoid variability of differentiation state and baseline contractility, ^[42]^ complicating data interpretation and functional analyses as well as large-scale, high-throughput screening ^[43–46]^.

To address these limitations, we developed a robust human iPSC-derived heart organoid model, termed “reCardioids”, designed to eliminate inter-organoid functional variability. By dissociating and reaggregating self-organizing cardioids, we achieved constructs with highly consistent rhythmic contractile dynamics and a more advanced state of cardiomyocyte maturation. Here, we comprehensively characterized the cellular and functional profile of reCardioids and validated their predictive utility by assessing their functional response to a clinically used anthracycline agent (doxorubicin) as well as characterizing both the functional and transcriptomic responses to ionizing γ-radiation.

## Results

### Formation of self-organizing cardioids with an outer lining of endothelial cells

In recent years, self-assembling heart organoids have become widely utilized in fields such as developmental research ^[25–27,29,30]^, tissue engineering ^[27]^ and disease modeling ^[25,47]^ among many others ^[42,48]^. Consequently, the diversity of heart organoids and the protocols to generate these have rapidly expanded. In parallel, there is growing interest in vascularizing tissues, including heart organoids, to improve their cardiac function and physiological recapitulation. However, studies detailing the vascularization of self-assembling heart organoids remain scarce^[25,26,28,30,49,50].^

Based on our review of existing studies exploring vascularization of self-assembling cardioids we hypothesized that the timing of vascular endothelial growth factor 165A (VEGF-165A; hereafter referred to as “VEGF”) addition during differentiation would play a role in vascular development. To generate heart organoids, we adapted a protocol by Hofbauer *et al.* describing the generation of cavity-containing heart organoids from hPSCs by controlling Activin, Bone morphogenetic protein (BMP), fibroblast growth factor (FGF), retinoic acid (RA) and Wnt pathways ^[26]^. First, this will generate mesodermal progenitors, which are then specified into cardiac mesoderm and finally cardiomyocyte progenitors. These organoids are termed cardioids and through addition of exogenous VEGF, they can be formed with an outer lining of endothelial cells. During the generation of cardioids with and without exogenous VEGF, we noticed that the timing of the VEGF addition in combination with the amount of iPSCs seeded, heavily influenced cardioid morphology and size (Figure 1A-C). As expected, seeding higher amounts of iPSCs (from 2,500 to 10,000 cells) resulted in larger cardioids, but also generally produced more consistently formed cardioids in terms of size, shape and function.

**Figure 1.**
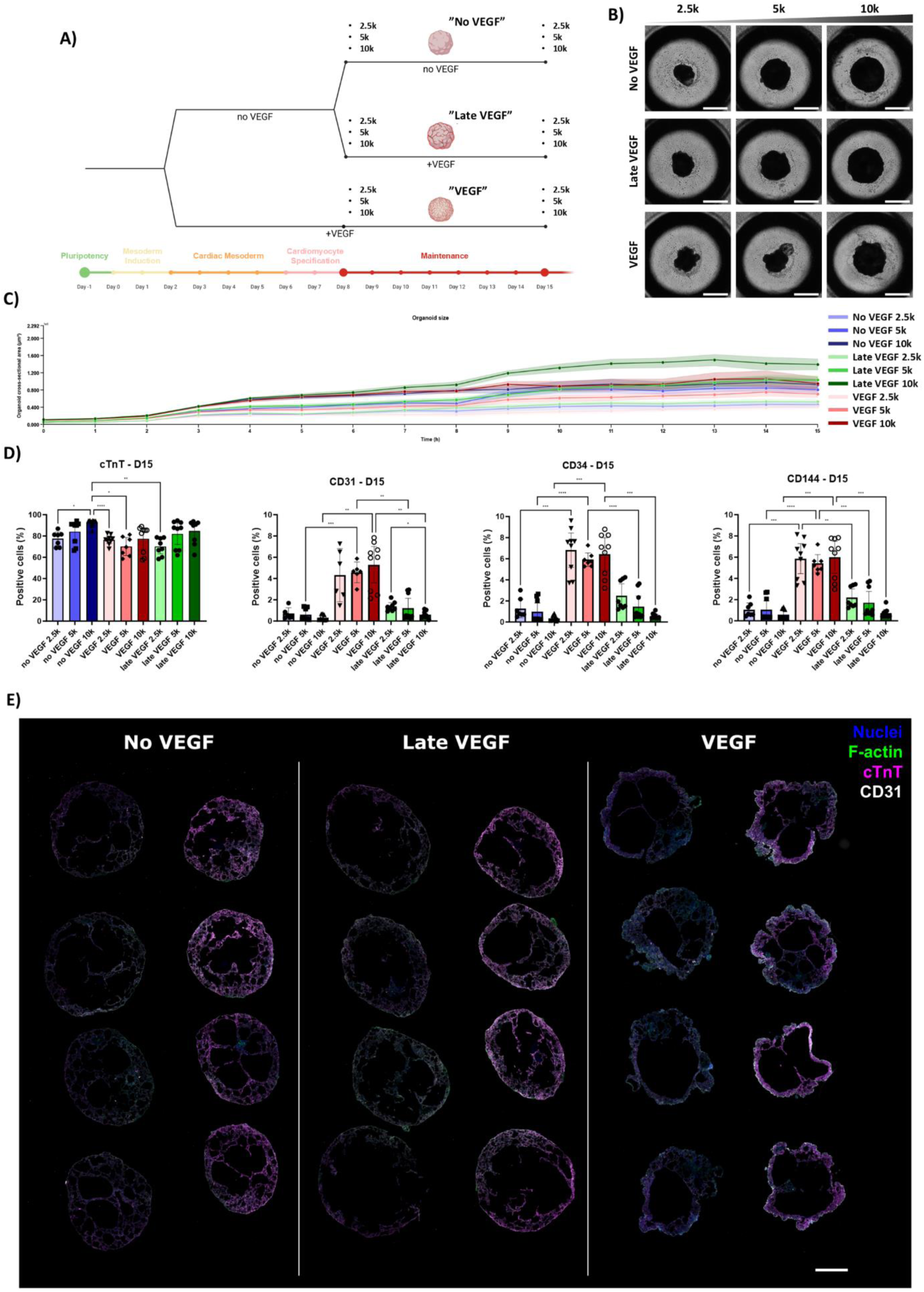
Cardioid starting iPSC amount and VEGF timing characterization. A) Schematic timeline of cardioid differentiation with indicated starting cells and VEGF condition. B) Brightfield images of all starting cell concentrations and VEGF conditions on day 15 of differentiation. Scale bar: 1 mm. C) Cardioid cross-sectional area over time during differentiation. Graphs show mean and 95% confidence interval. D) Quantification of flow cytometry data showing cell type-specific markers for cardiomyocytes and endothelial cells. Graphs show mean ± SD, * p<0.05 ** p<0.01 *** p<0.001 **** p<0.001, Kruskal-Wallis test, Dunn’s multiple comparisons test. E) Immunohistochemical staining of cryosectioned cardioids from each VEGF condition, generated using 10,000 starting iPSCs. Nuclei (blue), F-actin (green), cTnT) magenta), CD31 (white). Scale bar: 500 µm.

Regarding the timing of exogenous VEGF addition, three conditions were investigated (Figure 1A). VEGF was either (1) not added to the culture (hereafter referred to as the “no VEGF” condition), (2) added to the culture immediately before the cardiac mesoderm induction (hereafter referred to as the “VEGF” condition) or (3) added immediately after the cardiac mesoderm induction was completed (hereafter referred to as the “late VEGF” condition). When adding VEGF early, smaller protrusions were often formed on the surface of the cardioids (Figure 1B). On the other hand, when VEGF was not added or only added after cardiac mesoderm induction, the surface was typically smoother. From flow cytometric analysis of day 15 cardioids (Figure 1D), we found that all cardioids were composed mostly of cardiomyocytes (cardiac troponin T (cTnT)-positive cells), with the “no VEGF” cardioids formed from seeding 10,000 iPSCs, tending to contain the purest population of cardiomyocytes. Although both the “no VEGF” and “late VEGF” cardioids contained cells expressing endothelial markers (CD31+, CD34+, CD144+), the fraction of the endothelial population relative to the total cellular population was very low. In general, only the “VEGF” cardioids exhibited a significant population of endothelial cells. This was further validated by immunohistochemical analysis of cryosections, which also confirmed that these endothelial cells predominantly localized to the outer lining of the cardioids (Figure 1E, Figure S1, Figure S2). In addition, we were also able to confirm the presence of the fibroblast-like COL1A1+ cells also observed in Hofbauer *et al*. ^[26]^ (Figure S3, Figure S4).

### Dissociation and reaggregation improve the consistency of cardioid beating behavior

During routine maintenance of the cardioids, we observed considerable variability in their contraction rates. Typically, cardioids exhibited irregular fluctuations in peak-to-peak intervals, characterized by aperiodic or quasi-oscillatory patterns (Figure S5). This variability tended to increase with organoid size, and in the largest cardioids, contractions sometimes appeared as arrays of pulses rather than periodic contractions. We therefore investigated whether dissociating cardioids into single cells and subsequently reaggregating them at defined cell quantities could yield more consistent contractile behavior. To address this question, we generated a single-cell suspension of day 8 “VEGF” cardioids and reaggregated these at three different cell quantities: 25,000; 50,000 or 100,000 cells (A). This yielded organoids with uniform size while also enabling systematic modulation of their overall size (Figure 2C). We have termed these reaggregated constructs “reCardioids”.

**Figure 2.**
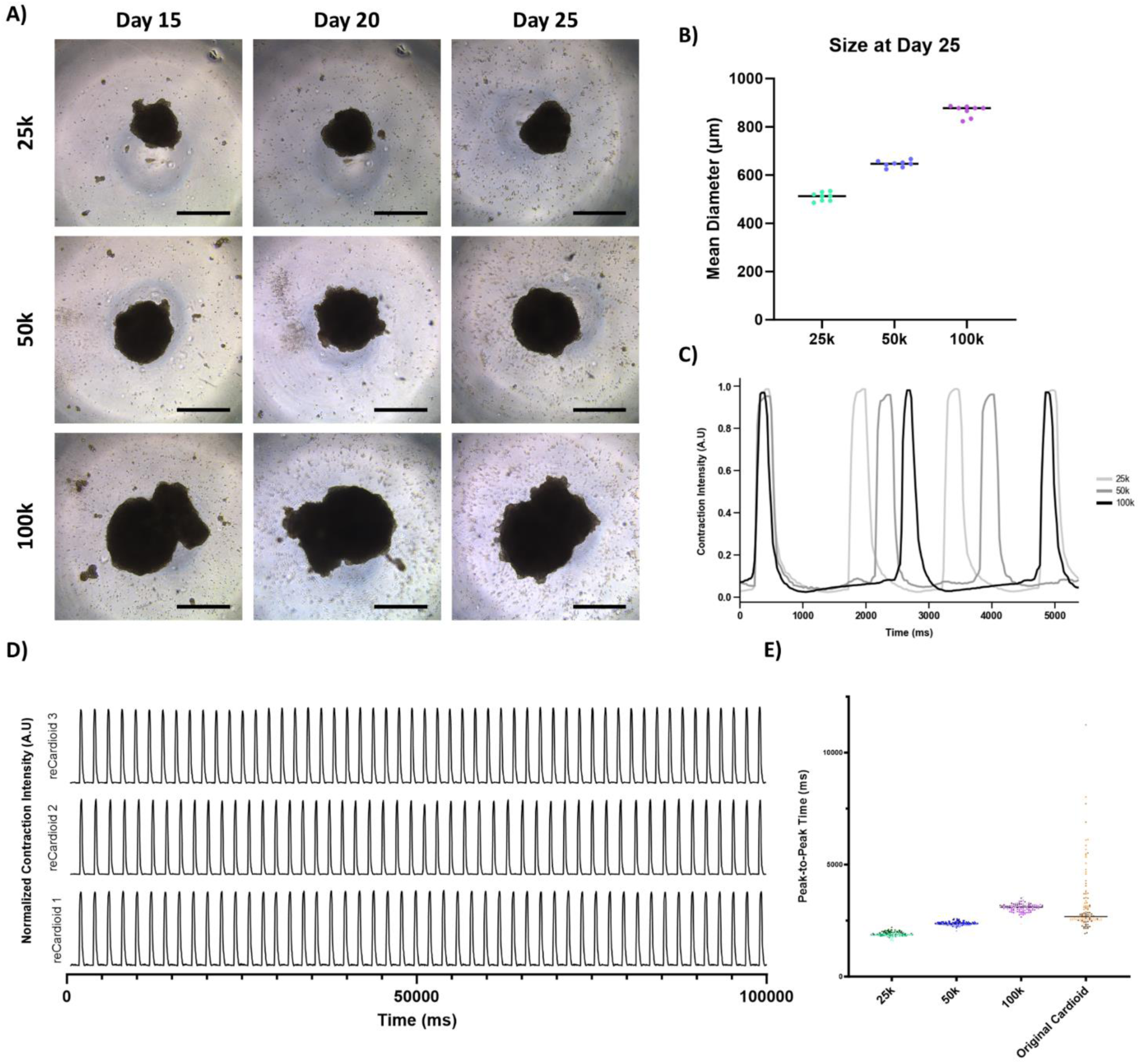
Generation of reCardioids. A) Brightfield images of reCardioids generated using either 25,000 (top), 50,000 (center) or 100,000 (bottom) cells during the first 25 days after the start of differentiation. Scale bar: 500 µm. B) Quantification of reCardioids diameters at day 25 after the start of differentiation. Mean ± SD. C) Representative contraction intensity profile of reCardioids generated using different numbers of cells, at day 25 after the start of differentiation. D) Representative traces of contraction of three individual reCardioids over 100 seconds, demonstrating consistent beating behaviour. E) Graph of individual peak-to-peak times of reCardioids and original cardioids (10,000 starting iPSCs) 25 days after the start of differentiation, n=3.

In general, reCardioids formed from 100,000 cells tended to develop into sheet-like aggregates, whereas those generated from 25,000 or 50,000 cells consistently formed more compact and uniformly sized structures. Notably, contraction frequency was highly consistent across all reCardioids and decreased with increasing total organoid size (Figure 2C & Figure S6). Importantly, in comparison to the original “VEGF” cardioids, age-matched reCardioids completely lacked irregular contraction behavior, instead displaying highly stable and rhythmic beating patterns (Figure 2D, 2E, Figure S6). Based on these results, we decided to continue with reCardioids formed using 50,000 cells .

### Single-cell RNA sequencing analysis of reCardioid composition

To further investigate the cellular composition of reCardioids and compare it with non-dissociated cardioids, we performed single-cell RNA sequencing (scRNA-seq) at days 15 and 25. Unsupervised clustering analysis revealed 16 distinct subpopulations (Figure 3A). Cell types were annotated based on classical cell type marker expression combined with CellTypist predictions. Nine of these clusters contained cells expressing established cardiomyocyte markers (clusters 4, 5, 6, 7, 9, 10, 11, 13, and 15). Notably, cluster 11 strongly expressed proliferative markers (Figure 3A, B, E). Consequently, we annotated cluster 11 as proliferative cardiomyocytes, while the remaining eight subsets were classified as general cardiomyocytes (CMs) (Figure 3A-D). In addition to the CM populations, clusters 3 and 14 contained cells expressing canonical endothelial cell markers, whereas cluster 2 exhibited a fibroblast-like transcriptional profile (Figure 3A, B, F, G, Figure S7). Together, these cell types represent the key lineages commonly identified in heart organoids, recapitulating the key features of early cardiac development. Interestingly, we also identified three distinct subsets (clusters 1, 12, and 16) defined by hepatic markers, including ALB, HNF4A, APOB, and APOA1 (Figure 3A, B, I-K). Unintended hepatic differentiation is a common off-target effect during the directed differentiation of iPSCs into cardiac lineages (and vice versa) due to shared induction and inhibition pathways [28,51,52]. This phenomenon reflects the close spatial proximity of the developing heart and foregut in the early human embryo, which exposes both progenitor populations to overlapping morphogen gradients and developmental timing. Lastly, we identified two clusters (cluster 0 and 8) that contained both cardiomyocyte- and hepatic-marker-expressing cells (Figure 3A-D, I-K). Unlike the other populations, these two clusters (cluster 0 and 8) did not segregate based on mature lineage identity. While the exact biological program driving their grouping remains undefined, they clearly share a distinct transcriptional state independent of lineage commitment. To better understand this population, we evaluated their marker expression and performed topology-based trajectory inference. Marker evaluation showed that these clusters were composed of a mixture of cells expressing different fetal lineage markers and showed no evidence of senescence (Figure 3A-D, I-K, Table S1). Trajectory network analysis demonstrated high connectivity specifically with both the committed cardiomyocyte and hepatic clusters, suggesting that these cells occupy an intermediate transcriptional state. We therefore annotated clusters 0 and 8 as mixed-lineage progenitors (Figure 3A, B, N-P). Analysis of cell type proportions revealed that CMs were the most abundant population at both timepoints across both protocols (reCardioids and non-dissociated cardioids). Furthermore, between days 15 and 25, the proportion of proliferating CMs and mixed-lineage progenitors decreased, while the overall CM population expanded in both protocols (Figure 3L, Figure S7A, D, E). This trend is consistent with ongoing tissue maturation and lineage commitment over time. At day 25, reCardioids exhibited a higher proportion of hepatic cells compared to non-dissociated cardioids, though the overall proportion of CMs remained comparable between the two groups (Figure 3L, Figure S7A, F). A closer examination of CM subpopulation dynamics at day 25 revealed protocol-specific differences. Cluster 5 was enriched in reCardioids, whereas clusters 4, 6, 7, 9, 10, and 13 constituted a larger fraction of the total CM pool in non-dissociated cardioids. Across these subsets, the most pronounced protocol-driven differences were observed in clusters 4, 5, and 10 (Figure 3M, Figure S7G, H, L). Longitudinal comparisons between day 15 and day 25 highlighted clusters 5, 6, and 10 as key populations associated with maturation. Notably, cluster 6 was proportionally the largest subset at day 15 but was substantially reduced by day 25. Conversely, clusters 5 and 10 comprised only a minimal fraction of total cells at day 15 but emerged prominently by day 25 (Figure 3M, Figure S7G, H, L). These two populations (clusters 5 and 10) also showed the greatest proportional variance between the two protocols. Looking further into these two clusters, we observed that while both subsets represent advanced stages of cardiomyocyte maturation, their transcriptional profiles suggest a divergence into specialized functional roles. Cluster 5 exhibited robust upregulation of genes essential for contractile force, mature sarcomere structure, and electrical stability, such as GJA1, DMD, MYH7, ATP2B1, and KCNJ2, pointing toward a mature working ventricular myocardium phenotype (Table S2). In contrast, cluster 10 displayed a marked enrichment of established conduction system and pacemaker-associated regulators, including ERBB4, TBX18, and HCN1, alongside comparatively lower expression of GJA1 (Table S2). To further investigate the broader transcriptomic shifts driven by the reCardioid protocol, we next compared the CM populations at day 25 using Gene Set Enrichment Analysis (GSEA). By evaluating Gene Ontology (GO) Biological Process (BP), Cellular Component (CC), and Molecular Function (MF) pathways, we aimed to define the overarching functional differences of the cells included in the CM group generated by each protocol. Among the top enriched terms, we observed a clear overrepresentation of pathways related to cardiac muscle formation and cardiac conduction. Specifically, reCardioids exhibited highly significant enrichment of pathways related to structural maturation, including “Contractile Muscle Fiber”, “Myofibril Assembly”, “I Band”, “M Band” and “Sarcomere Organization” (Figure 3Q). The structural advancement was accompanied by an increase in electrophysiological processes, evidenced by the enrichment of “Regulation of Heart Rate”, “Cardiac Muscle Cell Action Potential” and “Response to Calcium Ion” (Figure 3Q). Furthermore, reCardioids also showed a stronger enrichment of metabolic maturation pathways, including “Fatty Acid Beta-Oxidation” and “ATP Synthesis Coupled Electron Transport” (Figure 3Q). Additionally, enrichment of “Cytoplasmic Translation” and “Muscle Hypertrophy” pathways (Figure 3Q) suggested increased biosynthetic activity and cardiac muscle growth. Together, the enrichment of structural, electrical, and metabolic machinery suggests that the reCardioid protocol promotes a globally more mature cardiomyocyte phenotype. We also performed GSEA on the hepatic population and compared the enrichment of GO BP, CC, and MF terms of the hepatic cells at day 25 from both protocols. In reCardioids, the hepatic population showed enrichment of pathways predominantly related to advanced lipid metabolism, including “Triglyceride-Rich Lipoprotein Particle Remodeling” and “High-Density Lipoprotein Particle” networks, while actively downregulating immature “Glucose Catabolic Processes” (Figure 3R). This suggests that the hepatic population undergoes parallel functional maturation and may act as a localized metabolic hub, synthesizing and secreting the lipid substrates, helping to fuel the enriched fatty-acid beta-oxidation observed in the adjacent cardiomyocytes.

**Figure 3.**
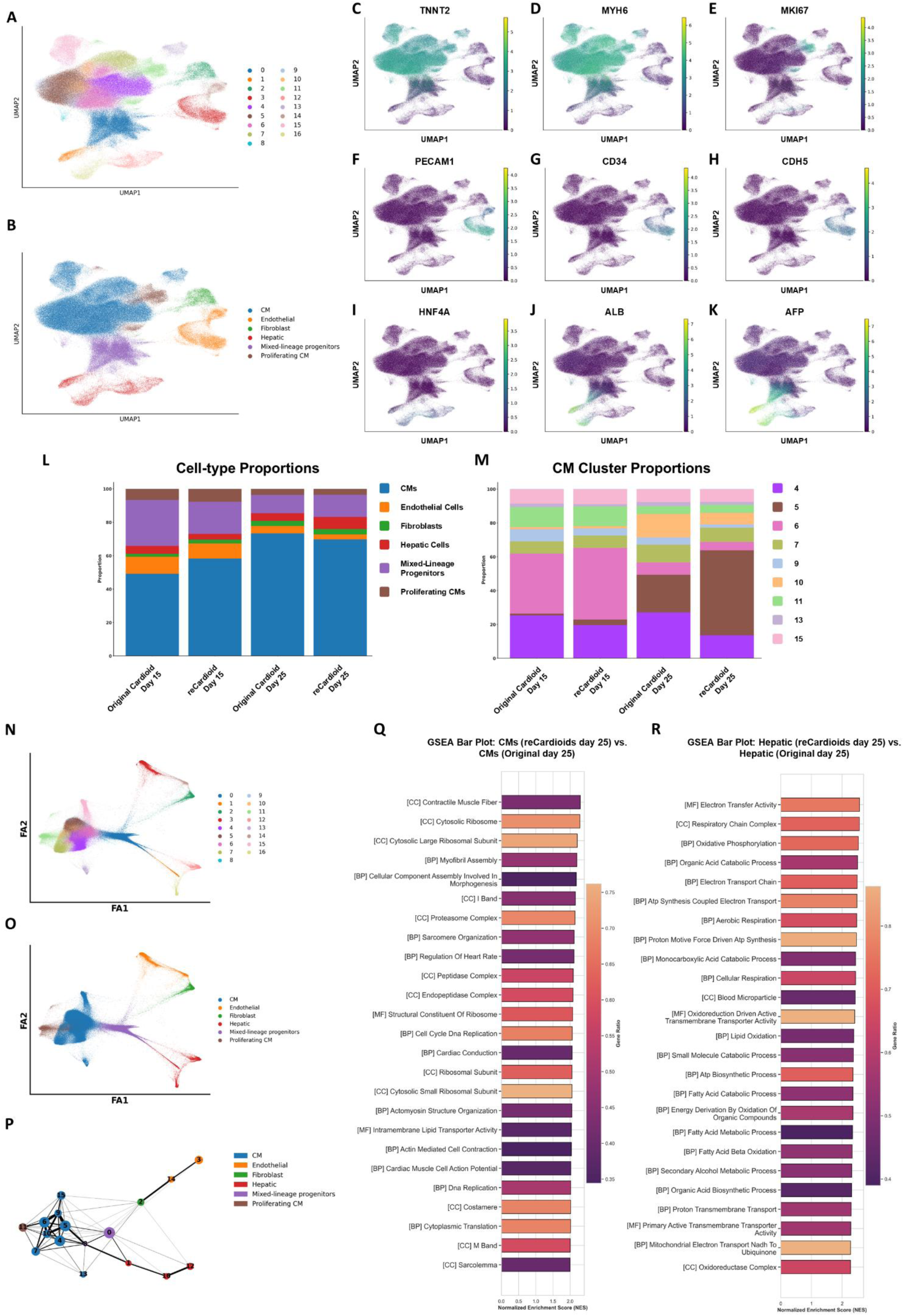
Single-cell transcriptomic profiling reveals cellular composition and enhanced cardiomyocyte maturation in reCardioids. (A) UMAP plot displaying 16 distinct cellular subpopulations identified through unsupervised clustering of cells from both protocols (original cardioids and reCardioids) at days 15 and 25. (B) UMAP plot displaying the manual annotation of major cell types based on canonical markers and CellTypist predictions, identifying cardiomyocytes (CM), proliferating CMs, endothelial cells, fibroblasts, hepatic cells, and mixed-lineage progenitors. (C-K) UMAP feature plots highlighting the expression of key lineage-specific markers: TNNT2 and MYH6 for cardiomyocytes, MKI67 for proliferating cells, PECAM1, CD34, and CDH5 for endothelial cells, and HNF4A, ALB, and AFP for hepatic lineages. (L-M) Stacked bar charts quantifying global cell-type proportions (L) and the proportional breakdown of individual cardiomyocyte subpopulations (M) across the different timepoints and protocols. (N-O) ForceAtlas2 (FA) embeddings of single cells initialized by PAGA, colored by the 16 unsupervised clusters (N) and major cell-type annotations (O) to visualize the continuous developmental progression. (P) Partition-based graph abstraction (PAGA) topology graph illustrating the inferred developmental trajectory and connectivity networks between the identified cellular clusters. Nodes represent individual clusters, and edge thicknesses denote the statistical measure of connectivity. (Q) GSEA bar plot comparing day 25 reCardioid CMs to original cardioid CMs. (R) GSEA bar plot for the hepatic subpopulation comparing day 25 reCardioid hepatic cells to original cardioid hepatic cells.

### ReCardioids reliably mimic doxorubicin-induced cardiotoxicity

To evaluate the ability of reCardioids to model clinically relevant cardiotoxicity, we exposed them to varying concentrations of doxorubicin, a widely used chemotherapeutic agent known for its dose-dependent cardiotoxic side effects. ReCardioids were treated with 0, 0.5, 1 or 5 µM doxorubicin for up to 72 h. At both 48 and 72 h, all doxorubicin-treated reCardioids exhibited a reduction in size compared to untreated controls (Figure 4A, B). In parallel, metabolic activity, assessed by total ATP production, declined in a dose- and time-dependent manner (Figure 4C), with higher doxorubicin concentrations leading to a more pronounced decrease. Together, the observed reductions in size and ATP levels suggest increased cell death following doxorubicin exposure. To confirm, we quantified lactate dehydrogenase (LDH) activity in the culture supernatant. As expected, LDH levels rose over time in all doxorubicin-treated conditions compared to controls, with the 5 µM group showing the highest activity (Figure 4D).

**Figure 4.**
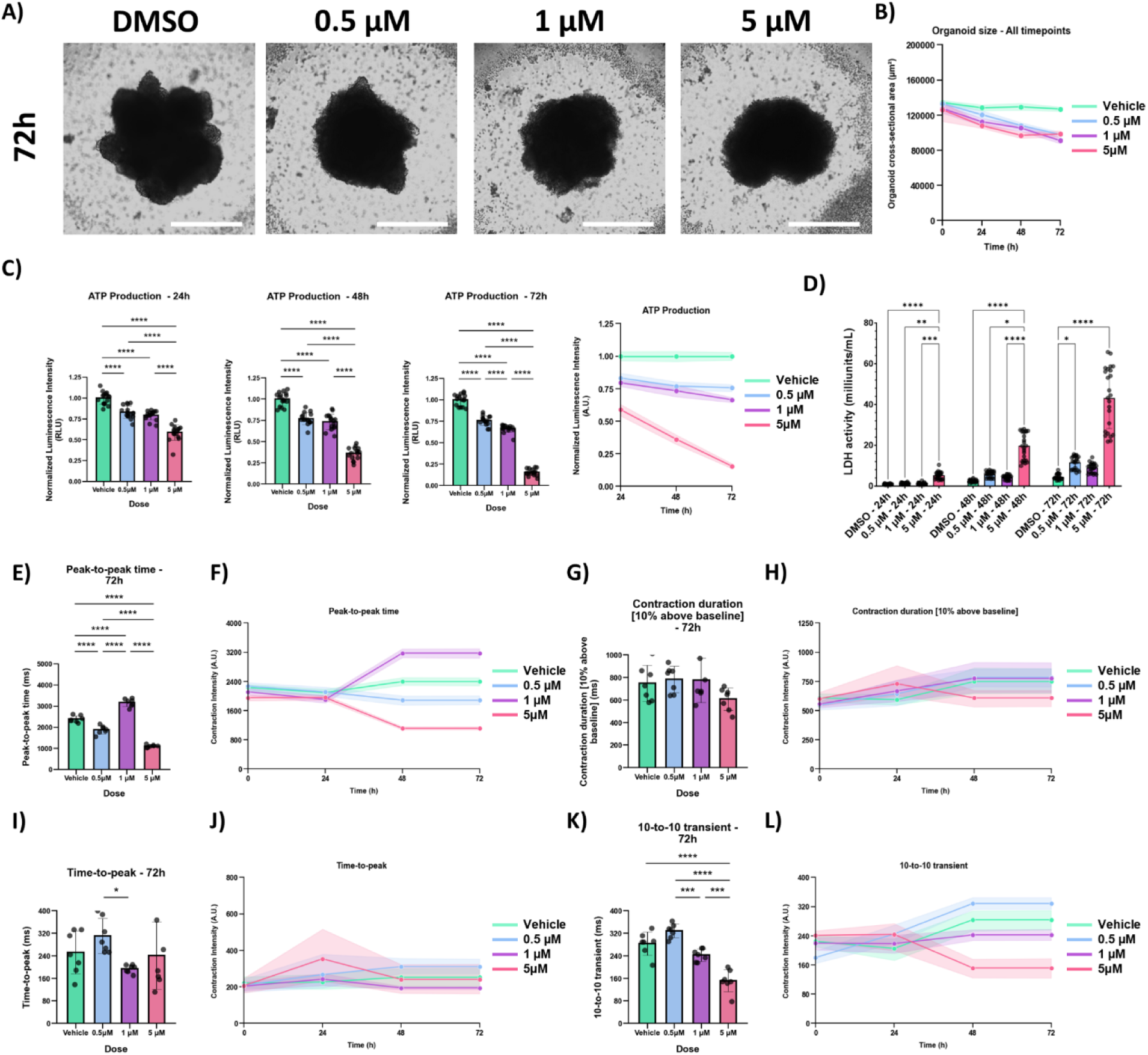
ReCardioids recapitulates cardiotoxic effect of chemotherapeutic drug doxorubicin. A) Brightfield images of reCardioids treated with doxorubicin for 72 h. Scale bar: 500 µm. B) Quantification of organoid size decline over time during exposure to doxorubicin. Graph showing mean ± 95% confidence interval. C) Quantification of ATP presence in reCardioids treated with doxorubicin for 24h, 48 h or 72 h. Bar plots: error bars shown as mean ± SD, lineplot: error bars shown as mean ± 95% confidence interval. D) Quantification of LDH release into supernatant media after 24 h, 48 h and 72 h doxorubicin exposure. Graph showing “accumulated” LDH activity at each timepoint in milliunits/mL of media. * p<0.05 ** p<0.01 *** p<0.001 **** p<0.001, Welch and Brown-Forsythe ANOVA. E-L) Quantification of spontaneous contraction parameters. E) Bar plot of peak-to-peak time after 72h of doxorubicin exposure. F) Line plot time course of peak-to-peak time from baseline to 72 h of doxorubicin treatment. G) Bar plot of contraction duration after 72 h of doxorubicin exposure. H) Line plot time course of contraction duration from baseline to 72 h of doxorubicin treatment. I) Bar plot of time-to-peak after 72 h of doxorubicin exposure. J) Line plot time course of time-to-peak from baseline to 72 h of doxorubicin treatment. K) Bar plot of 10-to-10 transient time (10% amplitude peak width) after 72 h of doxorubicin exposure. L) Line plot time course of 10-to-10 transient time (10% amplitude peak width) from baseline to 72 h of doxorubicin treatment. * p<0.05 ** p<0.01 *** p<0.001 **** p<0.001, Two-Way ANOVA with Tukey’s HSD Post Hoc Test.

Next, we assessed the contraction behavior of reCardioids following doxorubicin treatment. After 24 h, contraction parameters remained similar between treated and control groups (Figure 4E-L). However, by 48 and 72 h, significant differences in contraction frequency appeared across conditions (Figure 4E, F). Notably, the shift in contraction frequency at the 1 µM dose diverged from the overall dose-dependent trend observed across the other treatment groups. Additionally, we observed a dose-dependent reduction in the duration of the contracted state (Figure 4K, L), whereas time-to-peak (Figure 4I, J), relaxation duration, and the full contraction duration (Figure 4G, H) remained mostly unchanged.

### reCardioids as a model for radiotherapy-induced cardiotoxicity

Radiotherapy remains a cornerstone in the treatment of various cancers. However, when targeting tumors located in the anterior thoracic region, such as those in the lung or breast tissue, the heart is at risk of receiving unintended radiation exposure ^[12]^. Although modern techniques like proton therapy and 3D conformal radiotherapy have significantly reduced cardiac dose, the heart can still be subjected to a substantial radiation dose, particularly during photon-based lung cancer treatment ^[12]^. To assess the potential of reCardioids as a model for radiation-induced cardiotoxicity, we seeded them onto a fibrin matrix and exposed them to γ-rays from a Co-60 source, a standard source used in radiotherapy. ReCardioids were subjected to acute doses of 0.1, 0.5, or 1 Gy, alongside a 0 Gy sham control, and maintained in culture for up to 168 h post-irradiation (Figure 5A).

**Figure 5.**
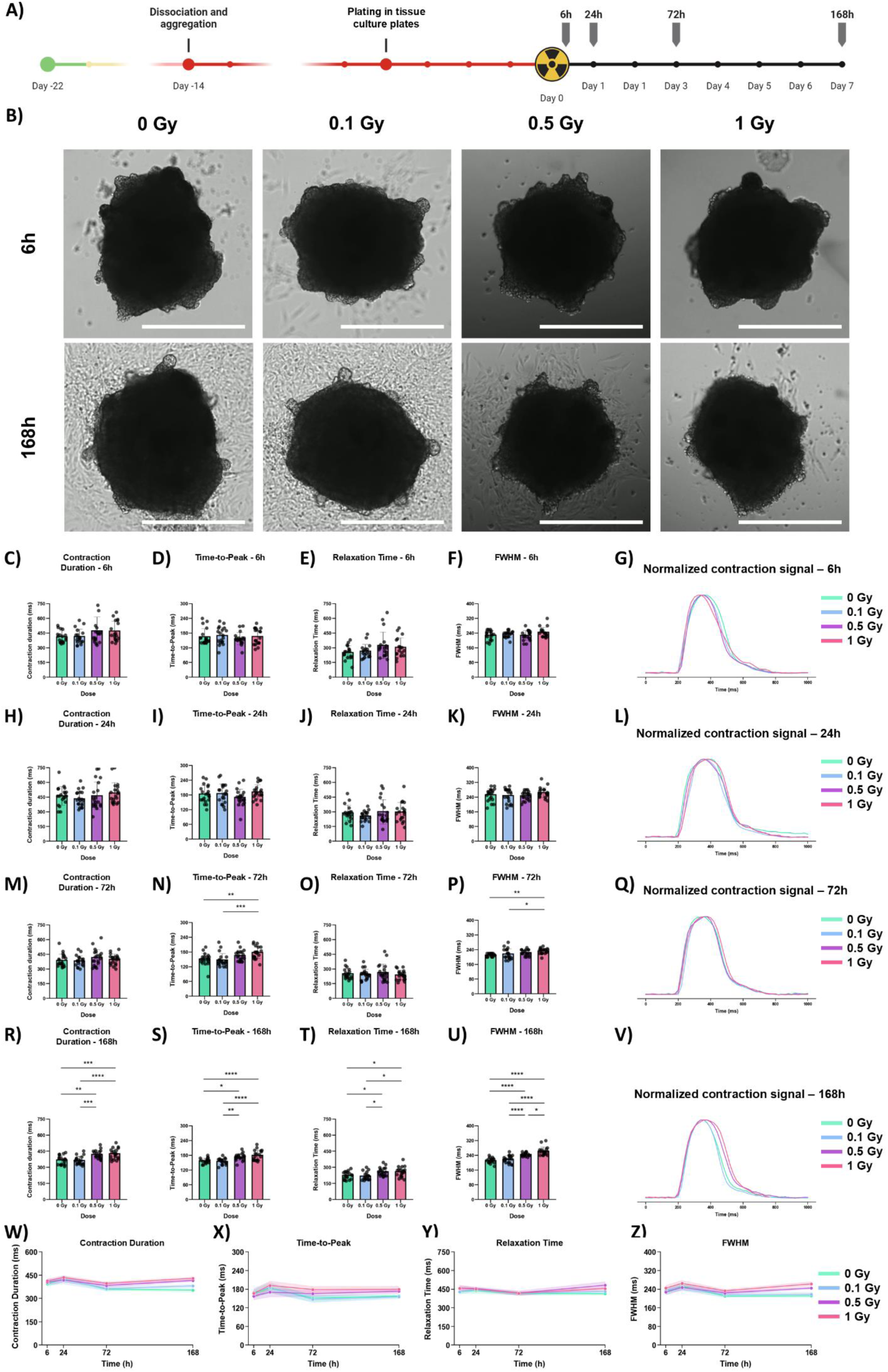
Exposure to radiotherapeutic ionizing radiation alters the contractile dynamics of reCardioids. A) Schematic timeline of experimental procedure. B) Brightfield images of irradiated and control reCardioids 6 h and 168 h post exposure to gamma-rays. Scale bar: 500 µm. C-F) Quantification of contraction duration (C), peak-to-peak time (D), relaxation time (E), time-to-peak (F) 6 h post exposure, error bars are shown as mean ± SD. G) Representative contraction profiles of reCardioids 6 h post exposure. H-L) Quantification of contraction duration (H), peak-to-peak time (I), relaxation time (J), time-to-peak (K) 24 h post exposure, error bars are shown as mean ± SD. L) Representative contraction profiles of reCardioids 24 h post exposure. M-Q) Quantification of contraction duration (M), peak-to-peak time (N), relaxation time (O), time-to-peak (P) 72 h post exposure, error bars are shown as mean ± SD. Q) Representative contraction profiles of reCardioids 72 h post exposure. R-V) Quantification of contraction duration (R), peak-to-peak time (S), relaxation time (T), time-to-peak (U) 168 h post exposure, error bars are shown as mean ± SD. V) Representative contraction profiles of reCardioids 168 h post exposure. W-Z) Line plot time course of contraction duration (W), time-to-peak (X), relaxation time (Y), FWHM (Z) from 6 h to 168 h. * p<0.05 ** p<0.01 *** p<0.001 **** p<0.001, Two-Way ANOVA with Tukey’s HSD Post Hoc Test

We first assessed the contraction behavior of reCardioids following irradiation (Figure 5B-Z). At both 6 and 24 h post-exposure, most contraction parameters remained largely unchanged across all groups, indicating minimal immediate effects (Figure 5C-L). However, by 72 h post-irradiation, reCardioids exposed to 1 Gy exhibited a significant increase in time-to-peak and full width at half maximum (FWHM), compared to the control group, along with several other altered metrics (Figure 5M-Q). At 168 h (7 days) post-irradiation, a dose-dependent trend became apparent across multiple contraction metrics (Figure 5R-V). While reCardioids exposed to 0.1 Gy showed similar contraction profiles to sham-treated controls, exposure to 0.5 Gy resulted in a significant prolongation of nearly all quantified contraction parameters relative to both the sham and 0.1 Gy groups (Figure 5R-V). Further increasing the dose to 1 Gy led to elevated values across nearly all contraction metrics compared to sham treated controls. Additionally, reCardioids irradiated with 1 Gy displayed significantly increased FWHM and 10-to-10 transient duration (time during full contraction) relative to the 0.5 Gy group (Figure 5R-V).

### Time-course clustering reveals a sequential trajectory from acute radiation response to pathological cardiac remodeling

To investigate the dynamic temporal transcriptional response of reCardioids following γ-ray exposure, we performed a Likelihood Ratio Test (LRT) to identify genes with significant expression changes over time and across doses, which yielded 1,412 differentially expressed genes (Figure 6A). To categorize their temporal patterns, we performed hierarchical clustering, which resolved seven distinct modules. For each module we conducted Gene ontology (GO) biological process overrepresentation analysis (Figure 6B) and calculated the first principal component (PC1) (Figure 6C), allowing us to track the biology involved in the module and the trajectory of its collective expression across all timepoints and doses. By integrating the functional profile of each module with their PC1 temporal profile, we identified a clear, dose-dependent timeline of the radiation response (Figure 6B, C).

**Figure 6.**
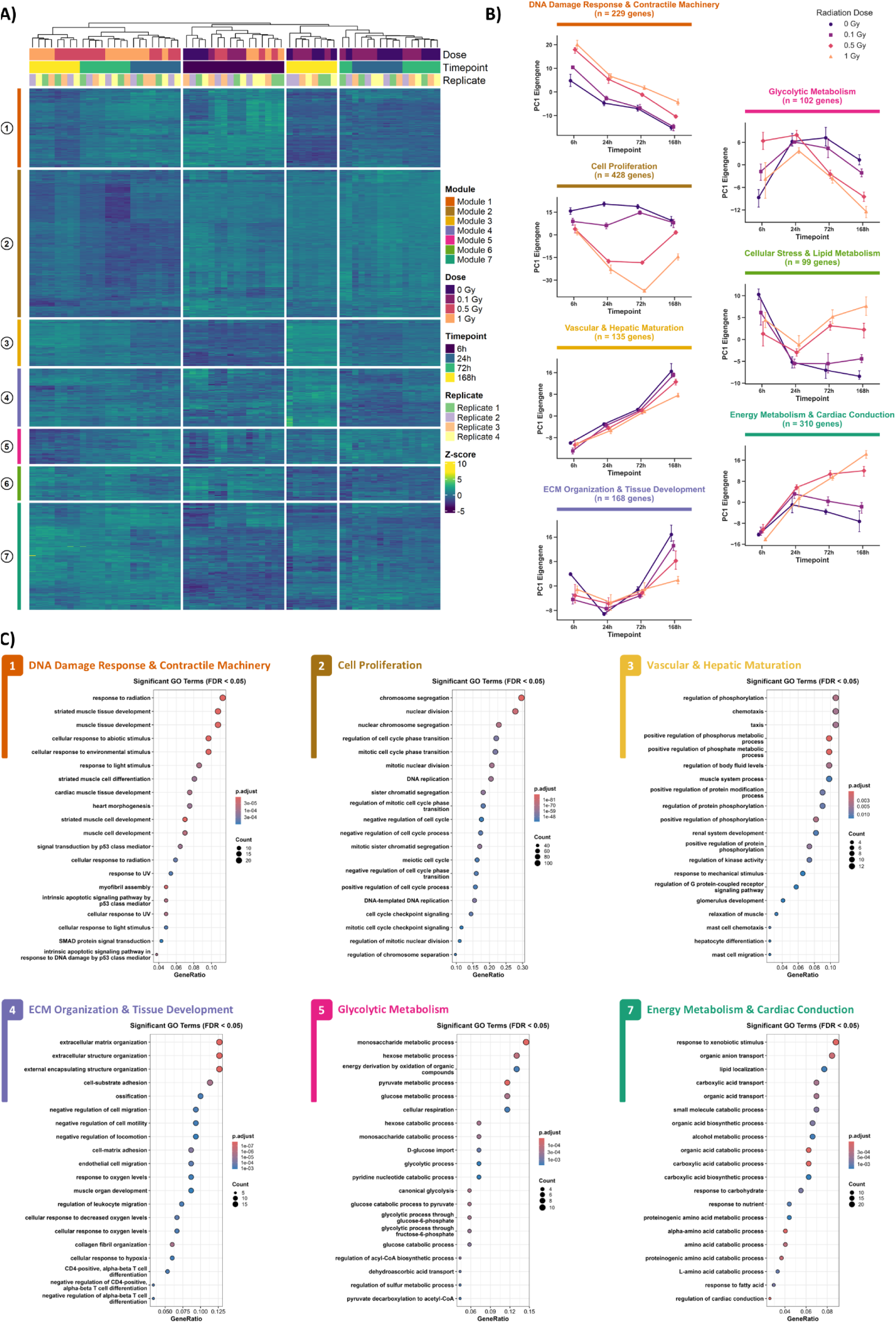
Bulk RNA sequencing reveals a dose-dependent temporal trajectory from acute radiation response to pathological remodeling. (A) Hierarchical clustering heatmap of 1,412 differentially expressed genes over a time course (6 h, 24 h, 72 h and 168 h) following exposure to varying doses of γ-radiation (0 Gy, 0.1 Gy, 0.5 Gy, and 1 Gy). Genes are partitioned into seven distinct temporal modules characterizing the evolving radiation response. (B) Line plots displaying the first principal component (PC1) trajectories for each of the seven modules. Data are presented as mean ± 95% CI. (C) Bubble plots illustrating Gene Ontology (GO) biological process overrepresentation for each module (FDR < 0.05).

The earliest transcriptional phase was captured in Module 1, DNA Damage Response & Contractile Machinery, where the PC1 profile exhibited a marked increase across all irradiated groups compared to the control at 6 h (Figure 6B). GO analysis revealed a classic, robust p53-driven DNA damage response, characterized by the upregulation of key regulatory genes (CDKN1A, MDM2 and SESN2). Following the initial 6 h peak, the response steadily declined, although it remained chronically elevated in a dose-dependent manner. By 168 h, only the 0.1 Gy group had returned to control levels (Figure 6B, C, Figure 7 and Table S3). Interestingly, in parallel with the DNA damage response, this module also contained a strong contribution from cardiac structural and contractile genes, such as TMOD1 and JPH2. Rather than reflecting decreased construction, this signature suggests an acute structural stress response and cytoskeletal remodeling ^[51–53]^ that stays chronically enriched in the 1 Gy group, as evidenced by an immediate upregulation of the contractile machinery that did not return to control levels (Figure 7, Table S3).

**Figure 7.**
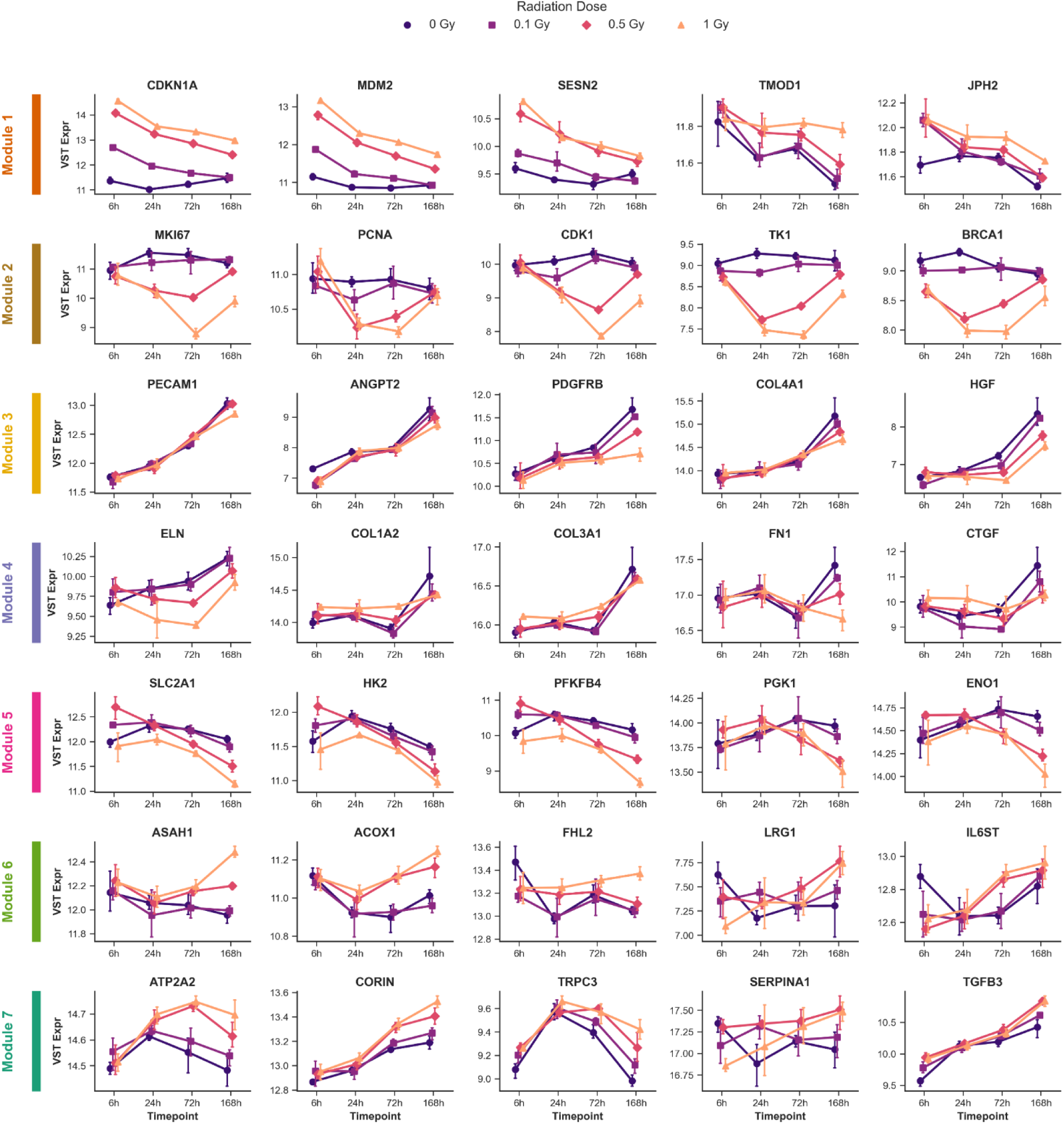
Temporal expression profiles of representative genes across identified transcriptional modules 1-7 (Module 1: DNA Damage Response & Contractile Machinery, Module 2: Cell Proliferation, Module 3: Vascular & Hepatic Maturation, Module 4: ECM Organization & Tissue Development, Module 5: Glycolytic Metabolism, Module 6: Cellular Stress & Lipid Metabolism, Module 7: Energy Metabolism & Cardiac Conduction). Line plots displaying the variance-stabilized transformed (VST) expression levels of key genes for the processes of interest in each module (Modules 1-7). Expression dynamics are tracked over a 168 h time course (6 h, 24 h, 72 h, and 168 h) following exposure to varying doses of ν-radiation (0 Gy, 0.1 Gy, 0.5 Gy, and 1 Gy). Data are presented as mean ± 95% CI.

The likely primary downstream consequence of p53 activation was observed in Module 2, Cell Proliferation, which was characterized by canonical markers of the cell cycle, mitosis and high-fidelity DNA repair (including MKI67, PCNA, CDK1 and TK1), along with GO terms covering cellular proliferation. This module exhibited a marked downregulation by 24 h in the irradiated groups and further decreased at the higher doses (0.5 Gy and 1 Gy) up to 72 h, after which they also maintained a lower expression at 168 h. Key proliferative markers followed the same trend, although in the 0.5 Gy group they mostly returned to control levels. This prolonged suppression (likely enforced by the acute p53/p21 response observed in Module 1, reflects a dose-dependent cell cycle arrest, leading to a significant but partially reversible disruption of the actively dividing progenitor pools (Figure 6B, C, Figure 7 and Table S3).

Starting at 72 h, two developmental modules capture a divergence between irradiated groups and the control group. Module 3, Vascular & Hepatic Maturation, represented the development of the organoid’s minority lineages, specifically vascular and hepatic cells. While the control group exhibited a steady upward trajectory, reflecting healthy maturation of these cellular compartments, irradiated organoids exhibited a deviating, impaired profile. This was evidenced by the presence of GO terms covering, vascular development, mural cell function and hepatic maturation. Correspondingly, irradiated organoids (primarily the 1 Gy group) exhibited decreased expression of endothelial and angiogenic markers (PECAM1, ANGPT2), vascular support cell markers (PDGFRB, COL4A1) and hepatic markers (HGF, CYP1A1), likely reflecting halted proliferation and dysfunction of these populations (Figure 6B, C, Figure 7 and Table S3). Simultaneously, Module 4, Extracellular Matrix (ECM) Organization & Tissue Development, revealed that the development of healthy ECM scaffold, expected during development of the tissue, was halted in the groups irradiated with 0.5 Gy or 1 Gy, as evidenced by the lower expression of mature matrix components like elastin (ELN). Although, notably, during the earlier phase (6-72 h), certain fibrillar collagens exhibited transient upregulation in the 0.5 Gy and 1 Gy groups, compared to the controls (Figure 6B, C, Figure 7 and Table S3).

The observed developmental stunting was followed by a decline in glycolytic metabolism, as illustrated by Module 5, Glycolytic Metabolism. While a decline in glycolytic metabolism is a normal feature in maturing cardiac tissues ^[54,55]^ (observed in 0 Gy controls), high dose exposure (0.5 Gy and 1 Gy) triggered a pronounced decline in glycolytic pathway components (including SLC2A1, HK2 and PFKFB4) of the organoids (Figure 6B, C, Figure 7 and Table S3).

In the final phase, the tissue underwent a state of pathological adaptation to cope with metabolic and structural failure. This was captured by two late-stage modules that diverged and surged upward exclusively in the irradiated samples. Module 6, Cellular Stress & Lipid Metabolism, lacked significantly overrepresented GO terms but was defined by a specific group of lipotoxic and biomechanical stress markers. As glycolysis declined, the tissue exhibited increased expression of lipid-clearing enzymes (*e.g.*, ASAH1, ACOX1), indicating the processing of toxic lipid byproducts, alongside mechanical stress sensors (*e.g.*, FHL2, LRG1) (Figure 6B, C, Figure 7 and Table S3).

Concurrently, Module 7 (Energy Metabolism & Cardiac Conduction) surged significantly at 168 h. This module was characterized by altered electrophysiological components (ATP2A2, SCN2B) and drivers of pathological hypertrophy and metabolic shifting (TRPC3, EGR1, ESRRG). Parallel to this module’s trajectory, we observed a concurrent trend toward dose-dependent elevation in classical clinical biomarkers of severe cardiac wall stress (NPPA, NPPB) ^[56,57]^ at 168 h in the highest dose groups, which was accompanied by the significant upregulation of their key activating enzyme, CORIN, within Module 7 ^[58,59]^. This continuous, compensatory upregulation of cardiac structural machinery (NEBL, NEXN) (similarly observed in Module 1) (Figure 6B, C, Figure 7 and Table S3) combined with the activation of these disease signatures, indicates that the tissue has ultimately undergone pathological, compensatory hypertrophic remodeling ^[60–62]^.

Furthermore, while classic interleukins associated with a typical Senescence-Associated Secretory Phenotype (SASP) were largely absent (Table S3), the high-dose irradiated groups (0.5 Gy and 1 Gy) exhibited elevated expression of specific tissue-protective and anti-proteolytic secretory proteins, including SERPINA1 and TIMP3. Notably, TGFB3 was also significantly upregulated. While TGF-β signaling is well-known to be involved in radiation-induced fibrosis ^[63]^, the specific elevation of the TGFB3 isoform (alongside SERPINA1 and TIMP3) (Figure 6B, C, Figure 7 and Table S3) has recently been shown to likely represent a complex tissue-repair response, mediating extracellular matrix remodeling and attempting to stabilize the surviving tissue following the acute damage. This interpretation is consistent with recent evidence that TGFB3 may preserve cardiac function and protect against unchecked fibrosis during heart failure ^[64,65]^.

## Discussion

Heart organoids have emerged as powerful tools for modeling human heart development, physiology, and disease ^[25–27,29,30,47]^. These 3D constructs derived from pluripotent stem cells offer a unique opportunity to recapitulate aspects of cardiac morphogenesis, cellular diversity, and functional behavior *in vitro*. Recent advances in self-organizing cardioids have significantly improved structural fidelity, enabling the formation of cavity-containing organoids with chamber-like features and spatial organization reminiscent of early heart development ^[26,27,30]^. However, despite these advances, challenges remain in achieving consistent contractile behavior. In this context, we developed reCardioids, reaggregated heart organoids with enhanced contractile regularity and reproducibility, to address these limitations and explore their utility in modeling cardiotoxicity induced by doxorubicin and thoracic radiotherapy.

At first, we aimed to optimize cardioids as a model for investigating cardiac responses to doxorubicin and radiation exposure. We selected cardioids due to their advanced tissue architecture, scalability and reproducibility. During early evaluations, however, we observed that while successful cardioid formation was reproducible, several parameters were not. In particular, their contraction behavior was highly variable from organoid to organoid. Individual organoids often exhibited irregular, quasi-oscillatory beating patterns, with inconsistent peak-to-peak intervals. This variability was especially evident in larger cardioids and progressively intensified with prolonged incubation without medium exchange. We speculate that this instability may be linked to a combination of conduction and metabolic factors, although further investigation is needed to confirm this hypothesis. More generally, contraction dynamics varied not only within individual organoids over time, but also between organoids produced under identical conditions. While this variability can be managed in high-replicate experiments, it becomes challenging when assessing subtle functional changes or sensitive readouts. In our experience, it also necessitates extremely frequent media changes to maintain stability, complicating experimental workflows. The reCardioid strategy addresses these challenges by reaggregating cardioid cells derived from pooled and dissociated cardioids from a single differentiation batch. This approach effectively eliminates inter-organoid variability stemming from slight differentiation heterogeneity and yields constructs with highly consistent contraction dynamics. Although reCardioids sacrifice some of the complex tissue architecture seen in cardioids, they offer distinct advantages, including improved functional consistency, tunable size and reduced experimental noise. These properties make reCardioids particularly well-suited for applications requiring precise quantification of contractile behavior, such as modeling heart rate variability or detecting subtle cardiotoxic effects.

To characterize the cellular differences between reCardioids and non-dissociated cardioids, we utilized scRNA-seq. Both protocols successfully generated the core cell types of the developing human heart, encompassing CMs, ECs, and fibroblast-like cells. The presence of these non-myocyte populations is crucial for the proper modeling of complex responses to environmental stress and injury, such as radiation-induced cardiotoxicity (where fibroblast and endothelial dysfunction play pivotal roles), as well as for drug screening ^[18,21,66]^. Our analysis also identified a distinct subpopulation of hepatic cells within both protocols. Unintended hepatic differentiation is a well-documented phenomenon in directed cardiac differentiation, driven by shared early morphogen programs and the developmental relationship between the cardiac and foregut lineages ^[28,67]^. While computationally separating this population was necessary to avoid confounding the CM transcriptomes, their presence may yield a functional advantage. The heart and liver are closely interconnected organs and their crosstalk is increasingly recognized as important in cardiometabolic diseases ^[68,69]^. Thus, co-developing hepatic cells may provide a more physiologically relevant model to study these systemic interactions ^[70]^. However, it is important to note that the precise anatomical configuration found *in vivo* is not recapitulated in this model.

Our concern when introducing a single-cell dissociation step into an established protocol is the risk of inducing significant shifts in cellular composition. Our data show that the overall cellular makeup remains highly comparable between the two protocols. While the proportional increase of hepatic cells in day 25 reCardioids could be viewed as an off-target effect, the core CM proportion was robustly maintained. More importantly, a detailed examination of the cardiomyocyte compartment revealed that the dissociation process neither negatively impacted cellular function nor stalled development. Furthermore, while the non-dissociated cardioids maintained a broader, more evenly distributed pool of CM subpopulations, the reCardioids exhibited a distinct enrichment for CMs with a more mature, working ventricular-like phenotype.

Building upon this phenotypic shift, *in vivo* cardiac maturation requires an extensive structural reorganization of the sarcomeres, coupled with an obligatory metabolic transition toward oxidative phosphorylation and advanced electrophysiological remodeling ^[54,55]^. Our transcriptomic analysis indicates that reCardioid CMs capture these developmental hallmarks to a greater extent than their non-dissociated counterparts. While the precise threshold of cardiomyocyte maturity required for accurate predictive modeling and toxicology remains a subject of ongoing debate ^[48,71]^, achieving a more advanced physiological baseline is universally recognized as highly advantageous for faithfully modeling complex cardiac responses ^[33,72]^.

Although reCardioids retain a predominantly fetal phenotype relative to adult tissue, the dissociation and reaggregation approach yields a comparatively more mature CM population. The precise mechanisms driving this effect remain to be fully elucidated, but the reaggregation of committed CMs may facilitate more efficient organization and structural integration. This spatial reorganization inherently increases cellular metabolic demand and contributes to the observed shift toward oxidative pathways ^[54,73]^. Furthermore, the highly stable and synchronous contractile rhythmicity observed in reCardioids leads to a more consistent cyclic strain on the developing muscle, providing well-established biophysical cues for cardiac growth and maturation ^[74,75]^. Intriguingly, this maturing CM profile parallels an advancement in lipid processing within the hepatic population. This points to a dynamic *in vitro* microenvironment where hepatic cells may act as a localized source of lipid substrates ^[68,72,76]^, actively sustaining the high energetic requirements of the maturing myocardium and supporting cardiac maturation.

Cardiotoxicity remains a significant concern in cancer therapy, with both chemotherapeutic agents and radiotherapy contributing to long-term cardiac dysfunction ^[78–82]^. Among these, doxorubicin notoriously causes cardiac dysfunction, with up to 9% of treated patients developing doxorubicin-induced cardiotoxicity ^[83,84]^. Emerging evidence suggests that, similarly to radiation-induced cardiotoxicity, doxorubicin may cause an accelerated aging-like phenotype, contributing to its clinical manifestations ^[7]^. Mechanistically and phenotypically, doxorubicin-induced cardiotoxicity shares striking similarities with radiation-induced cardiac injury ^[7,85]^. Based on this rationale, we used doxorubicin to validate the utility of reCardioids to model clinically relevant cardiotoxicity. Doxorubicin treatment led to reduced organoid size, decreased metabolic activity, elevated LDH release, and altered contraction behavior, effects that additionally became more pronounced with increasing dose and prolonged exposure. These findings are consistent with previous preclinical studies and, importantly, mirror the clinical observation that the severity of doxorubicin-induced cardiotoxicity correlates with cumulative dose. The unexpected shift (in contraction frequency) at 1 µM likely reflects a transitional phase of severe electro-mechanical disturbance. Overall, these results confirm that, similarly to other established advanced cardiac models ^[85–88]^, reCardioids can faithfully model key features of doxorubicin-induced cardiotoxicity.

To evaluate the ability of reCardioids to simulate the effects of clinically relevant radiation responses on the human heart, we next monitored the contractile performance of the organoids following γ-ray exposure. While baseline function was initially preserved, high-dose exposure (0.5 Gy and 1 Gy) resulted in significantly slower contraction dynamics by 168 h, compared to the control, consistent with the observed transcriptional trajectories.

The initial transcriptomic response confirmed the expected radiation response, capturing p53-driven DNA damage response (Module 1; Figure 6A-C) and a subsequent dose-dependent arrest of the dividing progenitor cells (Module 2; Figure 6A-C) ^[89,90]^. However, the broader value of the model is reflected in its ability to mimic certain aspects of the temporal pathogenesis of Radiation Induced Heart Disease (RIHD), beyond the acute DNA damage and proliferative arrest. Importantly, reCardioids captured multilineage deterioration of the irradiated heart, similar to the clinically recognized microvascular disease in RIHD ^[11,19,66]^. The reCardioids showed a dose-dependent impairment of vascular maturation (Module 3; Figure 6A-C), as evidenced by downregulation of endothelial and mural support markers such as PECAM1, ANGPT2, and PDGFRB, alongside key enrichment GO pathways such as “coronary vasculature development” (GO:0060976), “vascular endothelial growth factor signaling pathway” (GO:0038084), and “artery development” (GO:0060840). Since capillary rarefaction is a recognized event in clinical RIHD, these transcriptional signatures indicate that the model can recapitulate an important component of the disease process ^[11,19,66]^.

This vascular stunting was accompanied by metabolic alterations. While normal cardiac development involves a metabolic shift away from glycolysis ^[54,55]^, high-dose irradiation triggered an exaggerated decline of the glycolytic pathway (Module 5; Figure 6A-C). Consequently, the surviving tissue exhibited lipotoxic stress, as evidenced by the upregulation of lipid-clearing enzymes such as ACOX1 and ASAH1 (Module 6; Figure 6A-C). This transition recapitulates the loss of metabolic flexibility and the resulting lipotoxic environment characteristic of diseased or aging hearts ^[92–95]^.

Taken together, these transcriptional alterations (sustained DNA damage repair, proliferative arrest, vascular decline, and metabolic inflexibility) culminate in the observed functional deviation from the control (contraction dynamics) and a transcriptional signature of pathological hypertrophy (Module 7; Figure 6A-C). The late-stage alteration in electrophysiological machinery (ATP2A2/SERCA), together with a key regulator of metabolic remodeling (ESRRG) are hallmark indicators of a myocardium mounting a compensatory hypertrophic response ^[95–97]^. Concurrently, we observed a trend toward increased expression of classical clinical biomarkers of cardiac wall stress, NPPA and NPPB, ^[56,57]^ at 168 h in the highest dose group, alongside the significant dose-dependent upregulation of their activating enzyme, CORIN ^[58,59]^ (Module 7; Figure 6A-C). The parallel activation of remodeling drivers like TRPC3 indicates that the reCardioids execute a maladaptive hypertrophic program, resembling that seen in clinical RIHD ^[13,17,98]^.

Interestingly, we did not observe a classic inflammatory SASP after radiation exposure. Typically, higher doses of ionizing radiation induce a SASP in irradiated cells, driving pathological adaptations ^[66,99]^. Instead, the organoids here showed signs of mounting a sophisticated defense mechanism, where the late-stage upregulation of tissue-protective factors (notably TIMP3, SERPINA1, and TGFB3) suggests a complex attempt to stabilize damaged ECM and prevent proteolytic breakdown ^[64,65]^. It is important to note that while the current 168 h timeline successfully captures the initiation of pathological remodeling, longer-term studies will be necessary to observe the potential progression into end-stage disease. Collectively, this trajectory, progressing from microvascular stunting to metabolic crisis and hypertrophic remodeling, mirrors several aspects of *in vivo* pathogenesis of RIHD ^[19,100]^, offering an advanced human *in vitro* platform for further research.

In summary, reCardioids address the critical need for highly reproducible human heart organoids by combining contractile consistency with a more advanced cardiomyocyte maturity profile. By successfully recapitulating the complex pathogenesis of both doxorubicin and radiation-induced cardiac injury, this platform bridges the gap between simple *in vitro* assays and animal models. Ultimately, reCardioids provide a scalable and effective tool to study therapy-induced cardiotoxicity.

## Conclusion

As the population of cancer survivors increases, long-term, post-treatment quality of life is becoming an increasingly critical aspect of patient care. Consequently, understanding treatment-induced cardiotoxic effects, such as those caused by anthracyclines and radiotherapy, has grown increasingly urgent. Therefore, the development of highly accurate preclinical *in vitro* models to study these effects is essential. Here, we developed a robust heart organoid model, termed reCardioids, by dissociating and reaggregating self-assembling cardioids. These reCardioids exhibit low inter-organoid variability in size and contractile behavior while maintaining complex cellular composition. Furthermore, they demonstrate a more mature cardiomyocyte profile compared to their non-dissociated counterparts. Crucially, reCardioids can effectively simulate both doxorubicin- and radiotherapy-induced cardiotoxicity. They recapitulate key hallmarks of doxorubicin-induced damage, including decreased metabolism, cytotoxicity, and altered contractile dynamics. When exposed to radiation, the model successfully captures important aspects of radiation-induced heart disease. Bulk RNA sequencing revealed a temporal trajectory of injury following γ-ray exposure: beginning with a DNA damage response, progressing through vascular impairment and metabolic dysfunction, and ultimately resulting in compensatory pathological hypertrophic remodeling. In summary, this study demonstrates that the dissociation and reaggregation of self-assembling cardioids yields a more robust and mature platform for investigating common cardiotoxicities observed during cancer treatment.

## Materials and methods

### Cell culture and medium preparation

The human UCSD-165i-95-1 cell line was acquired from the WiCell Institute (University of California, San Diego, Dr. Kelly Frazer, Material Transfer Agreement Order Reference No. 5967, USA) ^[101,102]^. Cells were maintained under feeder free conditions according to the manufacturer’s instructions. Briefly, cells were grown as colonies on Matrigel (354277, Corning)-coated tissue culture plates in mTeSR Plus (STEMCELL Technologies, 100-0276). Medium was exchanged according to the manufacturer’s recommendations. Once colonies reached sufficient density (routinely after around 6-7 days), cells were passaged as clumps. PSC colonies were first detached using ReLeSR (STEMCELL Technologies, 100-0483) and broken apart into smaller clumps. Clumps were then seeded onto Matrigel-coated plates. Cells were tested negative for mycoplasma. Chemically defined medium (CDM) was prepared by mixing 50 % Iscove’s Modified Dulbecco’s Medium (IMDM, Gibco, 12440053) and F12 NUT-Mix (Gibco, 11765054) and adding bovine serum albumin (BSA, 5 mg/mL, Sigma-Aldrich, A7030), 1 % chemically defined concentrated lipids (Gibco, 11905031), 0.004 % 1-thiolglycerol (Sigma-Aldrich, M6145), and Transferrin (15 µg/mL, Sigma-Aldrich, 178481-100MG). The medium was then filtered using a 0.2 µm low-protein-binding filter.

### Human iPSC differentiation into cardioids

Cardioids were generated according to Hofbauer *et al.* ^[26]^. Briefly, iPSCs were cultured as colonies, harvested and counted. 2,500; 5,000 or 10,000 cells were seeded in a volume of 200 µl mTeSR Plus containing 5 mM Y-27632 (STEMCELL Technologies, 100-0276) in each well of an ultra-low attachment 96-well plate (Corning, 4520). Cells were collected at the bottom of the wells by centrifugation at 250 g for 5 min. Cells were left to form aggregates for 24 h, after which they were induced for 40-44 h with CDM medium containing FGF2 (30 ng/mL, STEMCELL Technologies, 78134), LY294002 (5 μM, STEMCELL Technologies, 72152), Activin A (50 ng/mL, STEMCELL Technologies 78132), BMP4 (10 ng/mL, BioTechne, 314-BP-010), and 6 µM CHIR99021 (STEMCELL Technologies, 72052) and 1 μg/mL of insulin (Roche, 11376497001). Then, the cells were induced with CDM medium containing BMP4 (10 ng/mL, BioTechne, 314-BP-010), FGF2 (8 ng/mL, STEMCELL Technologies, 78134), insulin (10 μg/mL, 11376497001), IWP2 (5 μM, STEMCELL Technologies) and retinoic acid (0.5 μM, Sigma Aldrich, #R2625) for 4 days while changing the medium every day. VEGF-165 (200 ng/mL, STEMCELL Technologies, 78159) was added to the culture where indicated. After 4 days the medium was changed to CDM medium containing BMP4 (10 ng/mL, BioTechne, 314-BP-010), FGF2 (8 ng/mL, STEMCELL Technologies, 78134), insulin (10 μg/mL, Roche, 11376497001) and VEGF-165 (100 ng/mL, STEMCELL Technologies, 78159), where indicated, for 2 days while changing the medium every day. Subsequently the cells were maintained in CDM medium containing insulin (10 μg/mL, Roche, 11376497001) and, if indicated, VEGF-165 (100 ng/mL, STEMCELL Technologies, 78159). The maintenance medium was exchanged every 2-3 days.

### VEGF-165 timing

VEGF-165 timing experiments were performed by differentiating 2,500; 5,000 or 10,000 starting iPSCs according to the previously detailed procedure. “No VEGF” conditions were differentiated without the addition of VEGF-165 at any of the steps. “VEGF” conditions were formed by the addition of VEGF-165 (200 ng/mL, STEMCELL Technologies, 78159) into the medium on day 3 for a total of 4 days, after which the differentiation proceeded with VEGF-165 as previously indicated. “Late VEGF” conditions were formed by adding VEGF-165 (100 ng/mL, STEMCELL Technologies, 78159) into the medium first on day 8 of differentiation and then maintained in VEGF-165-containing maintenance medium. Brightfield images were acquired every 24 h during differentiation and maturation using a Cytation 5 (BioTek) or Nikon Ti-E Eclipse inverted microscope equipped with a 5X objective. A custom-made Python script was used to automatically segment and determine the size of each cardioid. Cardioids were then fixed or dissociated for downstream analysis using immunofluorescent imaging and flow cytometry.

### Cryosectioning

Samples were fixed in 10% formalin (CellStor, BAF-6000-08A) overnight at 4°C. After fixation, samples were washed in 1X PBS and incubated overnight at 4°C in 30% (w/v) sucrose in PBS. The next day, the samples were embedded in TissueTek (Sakura, 4583) and frozen in 2-methylbutane (VWR, 24872) cooled to -60°C using dry ice. Samples were then stored at -20°C until sectioning. Sections were further stored at -20°C until subsequent processing.

### Immunofluorescence

Following fixation and sectioning, samples were left at room temperature for 15 min and then washed with 1X PBS three times for 5 min. Samples were then permeabilized using 0.1% Triton X-100 (Sigma Aldrich, T8532) in PBS for 15 min and then washed three times in 1X PBS for 5 min. Blocking was then performed using 5% goat serum (Thermo Fisher, 31873) TNB blocking solution (Perkin Elmer), prepared according to manufacturer’s instructions. Primary antibodies were then added to TNB containing 5% goat serum and incubated overnight at 4°C, after which the primary antibodies were washed by washing three times with 1X PBS for 5 min. Secondary antibodies were then added to TNB containing 5% goat serum at the indicated concentrations seen in Table 1. Antibody panel for immunofluorescent stainings and incubated at room temperature for 1-2 h. Slides were then washed three times with 1X PBS for 5 min and then stained with 4′,6-diamidino-2-phenylindole (DAPI, 5 µg/mL, Sigma Aldrich, D9542) in PBS. Where indicated, Alexa Fluor™ 488 or Alexa Fluor™ 594 Phalloidin (1:400, Invitrogen, A12379) was added to the DAPI solution to stain the f-actin.

**Table 1.** Antibody panel for immunofluorescent stainings.

| Antibody | Reference number | Manufacturer | Dilution |
| --- | --- | --- | --- |
| cTnT | ab209813 | Abcam | 1:294.7 |
| COL1A1 | 72026S | Cell signaling | 1:200 |
| CD31 | Ab24590 | Abcam | 1:100 |
| Goat anti rabbit | A11036 | Thermo scientific | 1:1000 |
| Goat anti mouse | A32733TR | Thermo scientific | 1:1000 |

### Cardioid dissociation

Cardioids were dissociated using STEMdiff™ cardiomyocyte dissociation kit (STEMCELL Technologies, 05025) according to the manufacturer’s instructions, with minor modifications. Briefly, cardioids were collected in a 1X PBS filled filter capped conical 15 mL tube. The cardioids were then washed twice in 1X PBS. The PBS was then removed and STEMdiff™ cardiomyocyte dissociation medium was added, after which the cardioids were incubated in an incubator at 37°C and with 5% CO_2_ for 10-18 min. Thereafter, STEMdiff™ cardiomyocyte support medium was added before spinning down the cells. Spun down cells were then resuspended in CDM medium containing insulin (10 μg/mL, Roche, 11376497001).

### Generation of reCardioids

Cardioids from the “VEGF” condition and generated from 10,000 iPSCs were dissociated according to the procedure described above. Unless otherwise indicated, 50,000 cardioid derived cells were then seeded in CDM medium containing insulin (10 μg/mL, Roche, 11376497001) and VEGF-165 (100 ng/mL, STEMCELL Technologies, 78159) inside each well of an ultra-low attachment 96-wellplate (Corning, 4520). Cells were collected at the bottom of the wells by centrifugation at 250 g for 5 min. Cells were left to form aggregates for 72 h. After 72 h the medium was changed, after which half the medium was changed every 2-3 days.

### Doxorubicin exposure

reCardioids were cultured until day 25 and were then treated with either a vehicle (dimethylsulfoxide, DMSO, Sigma Aldrich) or doxorubicin (Sigma Aldrich) at either 0.5 µM, 1 µM or 5 µM for 72 h. 100 µL supernatant was collected each 24 h and replaced with medium. Furthermore, each 24 h, videos of the contraction were acquired and the relative ATP production was quantified using the CellTiter-Glo® 3D Cell Viability Assay (Promega, G9682). Supernatant LDH activity was quantified using an LDH activity kit (Sigma Aldrich, MAK066).

### Irradiation of reCardioids

Irradiations were performed at the Laboratory of Nuclear Calibration (LNK) of the Belgian Nuclear Research Center (SCK CEN) in Mol, Belgium, an ISO 17025:2017 accredited facility. reCardioids were seeded in fibrinogen coated 96-well plates at day 21. reCardioids were then irradiated vertically in free air by γ-rays using a Co-60 γ-ray source (1.1732, 1.3325 MeV) with a 35 × 35 cm collimated beam at a distance of 100 cm. Dosimetry was performed in accordance with ISO 4037:2019, with air kerma (K_a_) traceable to the LNK primary standard graphite cavity chamber ^[102]^. Embedded reCardioids were irradiated with a total dose of either 0 Gy (sham treated), 0.1 Gy, 0.5 Gy or 1 Gy using a dose rate of approximately 0.66 Gy/min. Sham treated reCardioids were kept at room temperature (21-22 °C) during the irradiation. Videos of embedded reCardioids were then acquired 6 h, 24 h, 72 h or 168 h after irradiation. Furthermore, embedded reCardioids were either snapfrozen or fixed at the same time points for downstream analysis.

### Contraction analysis

Live imaging was performed within a heated chamber (OKO Labs) at 37°C using a Nikon Eclipse Ti-E inverted microscope equipped with a 5X objective. Uncompressed live videos of tissues were captured using an exposure time of 100 µs at 45-50 FPS. Videos were cropped and the contraction was evaluated using the MUSCLEMOTION ^[103]^ Fiji plug in. Videos used for long term contraction profiles were cropped and evaluated using a custom Python script. Briefly, movement magnitude was quantified via Farneback’s dense optical flow algorithm utilizing a reference frame that evolved gradually between contractions and was fixed immediately prior to each contraction event.

### Flow cytometry

After dissociation, cardioid derived cells were placed on ice and counted. Cells were then stained with LIVE/DEAD™ Fixable Violet Dead Cell Stain Kit (Invitrogen, L34963) according to the manufacturer’s instructions. After staining, cells were washed with 1X PBS and then stained with a multicolor immunofluorescent panel found in Table 2 Samples were then run through a Novocyte Quanteon for data acquisition.

**Table 2.** Flow cytometry antibody panel.

| Protein | Fluorophore | Reference number | Manufacturer | Dilution |
| --- | --- | --- | --- | --- |
| CD31 | FITC | 303103 | Biolegend | 5 μL/(million cells) in 100 μL staining volume |
| CD34 | PE | 343605 | Biolegend | 5 μL/(million cells) in 100 μL staining volume |
| CD144 | BV786 | 565762 | BD Biosciences | 5 μL/(million cells) in 100 μL staining volume |
| cTnT | Alexa™ 647 | 565744 | BD Biosciences | 5 μL/(million cells) in 100 μL staining volume |

### Image acquisition

Fluorescent images were acquired using a Nikon Eclipse Ti-E inverted microscope, equipped with a 20X objective using a Teledyne BSI Prime camera. Live imaging was performed inside of a heated chamber (OKO Labs) at 37°C using the same Nikon Eclipse Ti-E inverted microscope, equipped with a 5X objective. Videos and images were captured using a Teledyne Prime BSI camera.

### Cell preparation for scRNA-seq

On day 15 and day 25, Cardioids (VEGF condition, formed from 10,000 iPSCs) and reCardioids were dissociated and fixed for scRNA-seq. Briefly, two cardioids or three reCardioids were pooled per sample, to acquire sufficient cells for the Evercode Low Input Cell and Nuclei Fixation v3 kits (PARSE Biosciences). Cardioids and reCardioids were then dissociated using the STEMdiff™ cardiomyocyte dissociation kit (STEMCELL Technologies), according to the manufacturer’s direction with small modifications. In short, organoids were washed three times using 1X PBS, after which they were incubated in STEMdiff™ cardiomyocyte dissociation medium for 7 min inside an incubator (37°C, 5% CO_2_). After incubation, the dissociation medium was removed and pipetted into ice-cold STEMdiff™ cardiomyocyte support medium. Pre-warmed dissociation medium was then pipetted onto the organoids and then incubated for 5 min inside an incubator. After incubation, the dissociation medium was removed and pipetted into the ice-cold support medium and the process was repeated. For day 15 samples, 5 incubations were performed in total, while 6 incubations were performed for day 25 samples. Cells were then centrifuged (5 min, 200 g), support medium was aspirated and the cells were resuspended in CDM medium containing insulin (10 μg/mL, Roche, 11376497001) and VEGF-165 (100 ng/mL, STEMCELL Technologies, 78159). Afterwards, cells were counted and 100,000 cells were collected and fixed using the Evercode Low Input Cell and Nuclei Fixation v3, according to the manufacturer’s instructions. Samples were then stored at -80°C until barcoding and library preparation.

### scRNA sequencing

Cells were thawed, recovered and counted according to the manufacturer’s instructions of the Evercode Low Input Cell and Nuclei Fixation v3 kits. Barcoding and library preparation were then performed using the Evercode WT Mega v3 kit (PARSE Biosciences). For each sample, 10,000 cells were barcoded, pooled and then divided into 16 sublibraries. Lysis, cDNA amplification and sequencing library preparation were then performed on these 16 sublibraries. Libraries were sequenced by PARSE Biosciences using an Ultima UG 100 sequencing platform and data was preprocessed using the pipeline in the Trailmaker platform. Bioinformatics analysis of the processed data was performed with Python (version 3.12.12). Scanpy (version 1.12) was used for quality control, dimension reduction, clustering and trajectory analysis. Cell types were manually assigned to resulting clusters using marker genes as well as using celltypist (version 1.7.1) labels. Differential expression analysis was performed using pydeseq2 (version 0.5.4) and characterization of the resulting gene lists were performed with gseapy (version 1.1.11). Compositional analysis was performed with scCODA (version 0.1.9).

### Bulk RNA sequencing

RNA was extracted using TriZol reagent (Invitrogen) and DNA was removed using TURBO DNA-free kit (Invitrogen), according to the manufacturer’s instructions. Sample preparation for sequencing was performed by Novogene (Münich, Germany). Messenger RNA was purified from total RNA using poly-T oligo-attached magnetic beads. After fragmentation, the first strand cDNA was synthesized using random hexamer primers, followed by the second strand cDNA synthesis using dTTP, end repair, A-tailing, adapter ligation, size selection, amplification, and purification. Sequencing was performed using an Illumina NovaSeq X.

Resulting reads were mapped to hg38 using hisat (version 2.2.1) and quantified using featureCounts (version 2.0.6). Differential expression analysis was performed with DESeq2 (version 1.42.0) and functional characterization of the differentially expressed genes were performed with the R package clusterProfiler (version 4.18.4); these were performed with R (version 4.5.2).

### Statistics

Quantitative data are expressed as the mean ± standard deviation (SD) or the mean ± 95% confidence interval (CI), as indicated in the figure legends. Statistical significance was defined as *p* < 0.05, with specific thresholds denoted by asterisks (* *p* < 0.05, ** *p* < 0.01, *** *p* < 0.001, **** *p* < 0.0001).

Depending on the experimental design, the following statistical tests were employed:

- Flow Cytometry: Differences in cell type-specific markers were evaluated using the Kruskal-Wallis test followed by Dunn’s multiple comparisons test.
- Cardiotoxicity Assays: Accumulated LDH activity was analyzed using Welch and Brown-Forsythe ANOVA tests.
- Contraction Dynamics: Changes in spontaneous contraction parameters (e.g., peak-to-peak time, contraction duration, time-to-peak, relaxation time, and FWHM) were assessed using a Two-Way ANOVA with Tukey’s Honestly Significant Difference (HSD) post-hoc test.
- Transcriptomic Analysis (Bulk RNA-Seq): Differential gene expression testing for time-dose interactions was performed using a Likelihood Ratio Test (LRT). Significance for differential expression was determined using a Benjamini-Hochberg adjusted *p*-value < 0.05. For Gene Ontology (GO) biological process overrepresentation analysis, statistical significance was defined by a False Discovery Rate (FDR) < 0.05.

## Acknowledgements

We thank the Cell Press editorial team and reviewers for their insightful feedback. We also acknowledge the members of the Astrocardia and PULSE consortium for their insightful discussions and scientific feedback. In particularly the BIO INX team for their valuable insights regarding the versatility, application potential, and technical requirements associated with the development of the heart organoid. We also thank the QbD Group for their support in quality verification and for their contributions toward establishing an improved and robust workflow for organoid generation and validation. Additionally, we thank Gustavo Tarqueto Duarte and Serena Bordignon from the Biosphere Impact Studies group at SCK CEN for their insightful feedback and help regarding the RNA extraction. Lastly, the authors thank Bart Marlein, Raf Aarts and Cristian Mihailescu from the Dosimetric and Calibration Services group at SCK CEN for help performing the irradiations.

## Contributors

E.R., B.B. and K.T. conceptualized the study. E.R., B.B., Z.J., B.C., C.V.R. and R.V. performed the experimental work. E.E. and E.R. performed and analysed the RNA sequencing data. All the authors contributed to the writing, reviewing and editing of the manuscript. K.T., S.B, L.M. acquired the funding. All the authors have read and approved the final manuscript.

## Data availability statement

The scRNA-seq data that supports the findings of this study have been deposited in ArrayExpress under accession number E-MTAB-16965 and the bulk RNA-seq data have been deposited in ArrayExpress under accession number E-MTAB-16889. Summary and processed data supporting the findings of this study are included in the article and its Supplementary Information.

## Supplementary material

Table S1. List of top differentially expressed genes for the Mixed-lineage progenitor cells compared to all CMs, all Hepatic cells or all non-CM and non-Hepatic cells of the scRNA-seq data.

Table S2. List of top differentially expressed genes for each CM cluster compared to all other CM clusters from the unsupervised clustering of scRNA-seq data.

Table S3. List of all genes identified by LRT analysis.

**Figure S1.**
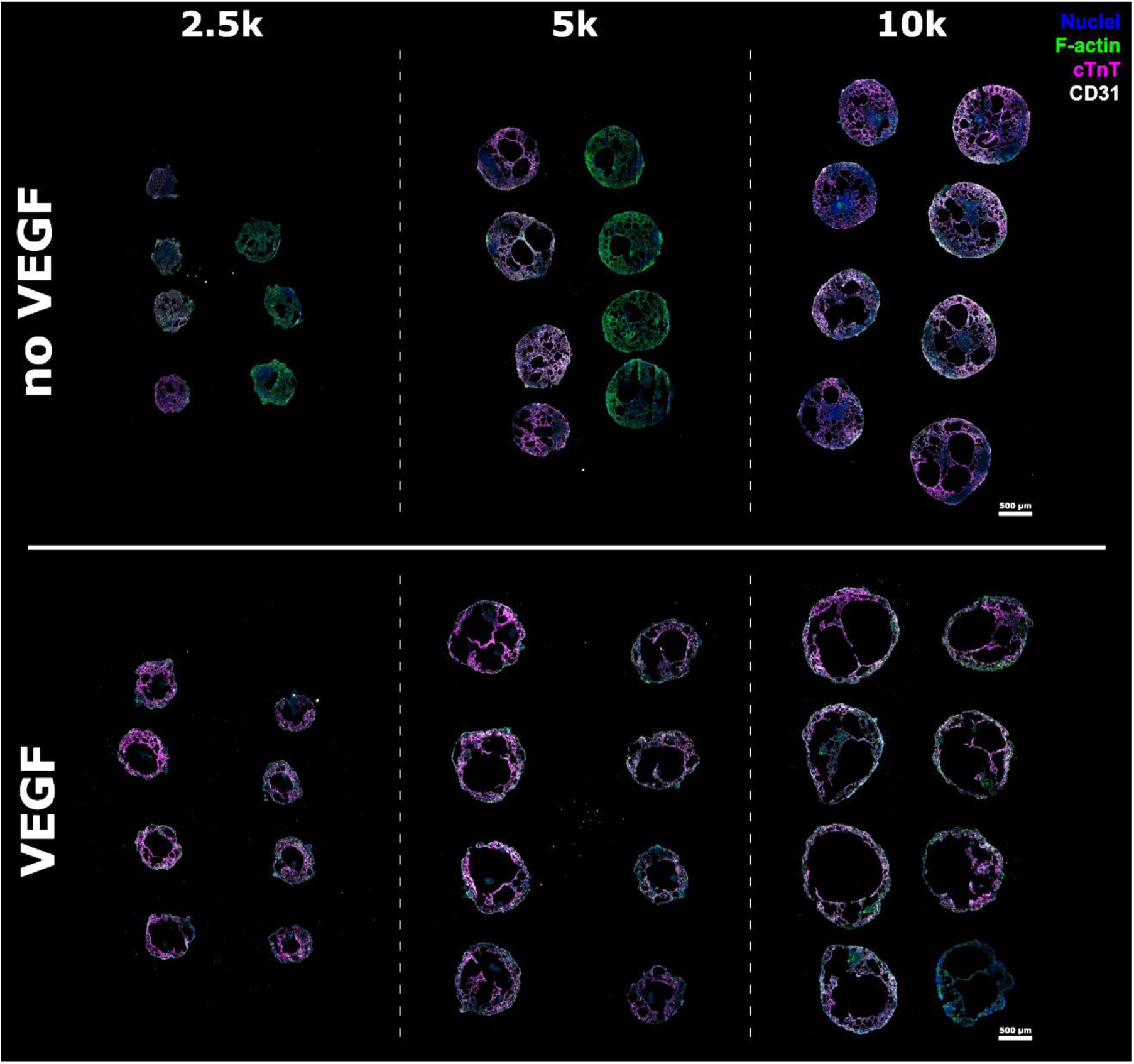
Immunohistochemical staining of cryosectioned cardioids (day 8) from each VEGF condition, generated using 2,500; 5,000 or 10,000 starting iPSCs. Nuclei (blue), F-actin (green), cTnT (magenta), CD31 (white). Scale bar: 500 µm.

**Figure S2.**
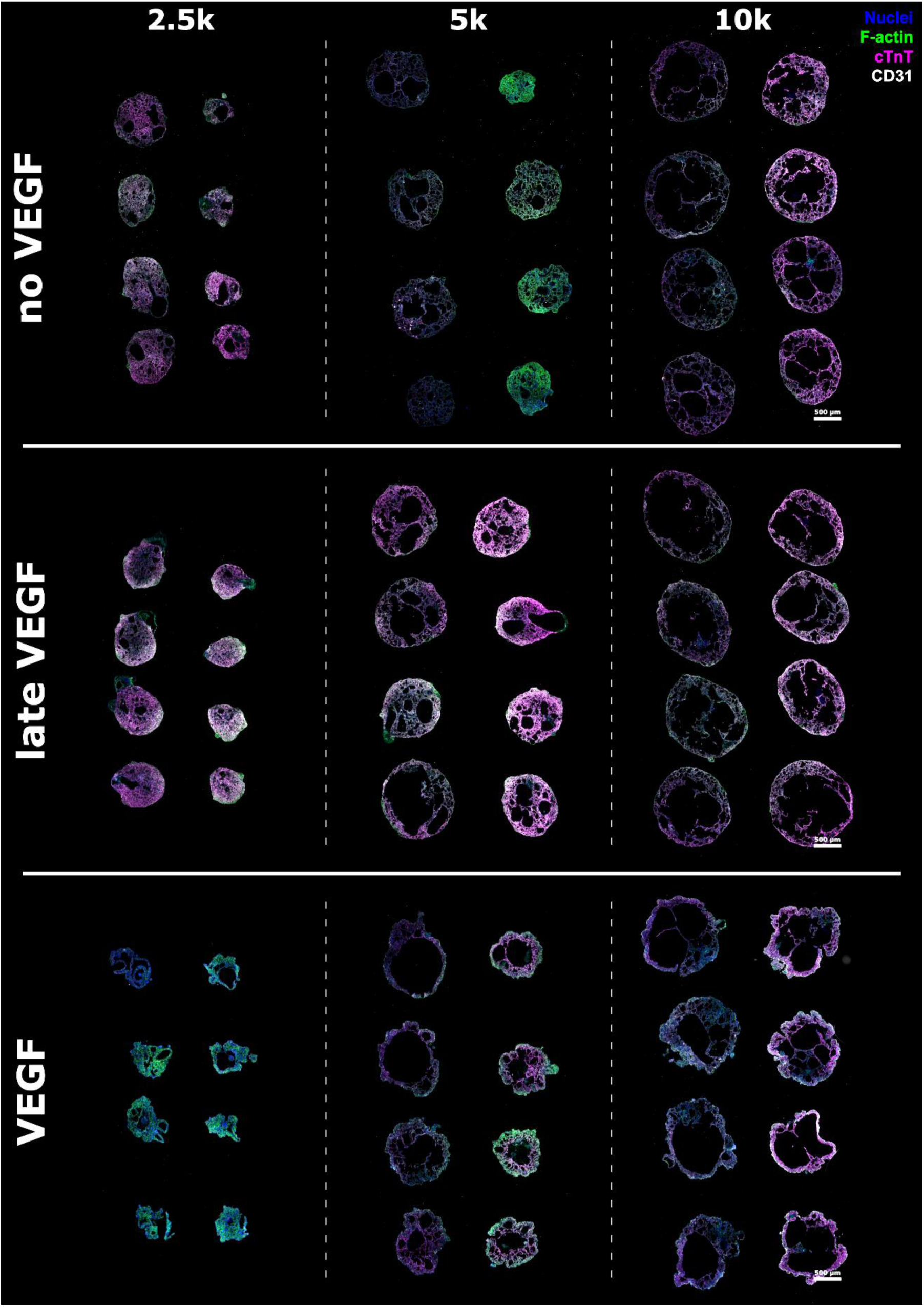
Immunohistochemical staining of cryosectioned cardioids (day 15) from each VEGF condition, generated using 2,500; 5,000 or 10,000 starting iPSCs. Nuclei (blue), F-actin (green), cTnT (magenta), CD31 (white). Scale bar: 500 µm.

**Figure S3.**
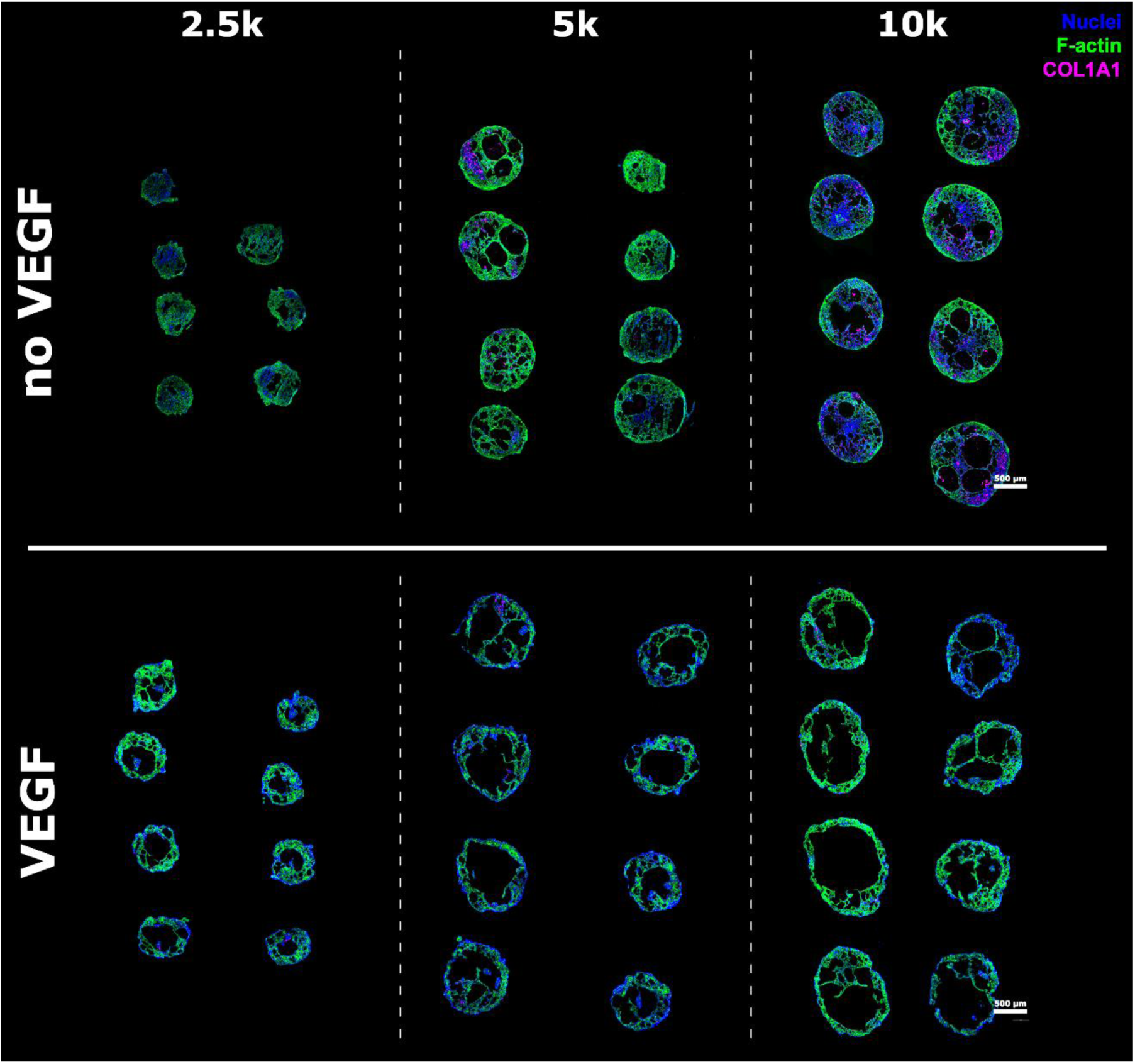
Immunohistochemical staining of cryosectioned cardioids (day 8) from each VEGF condition, generated using 2,500; 5,000 or 10,000 starting iPSCs. Nuclei (blue), F-actin (green), cTnT (magenta), COL1A1 (white). Scale bar: 500 µm.

**Figure S4.**
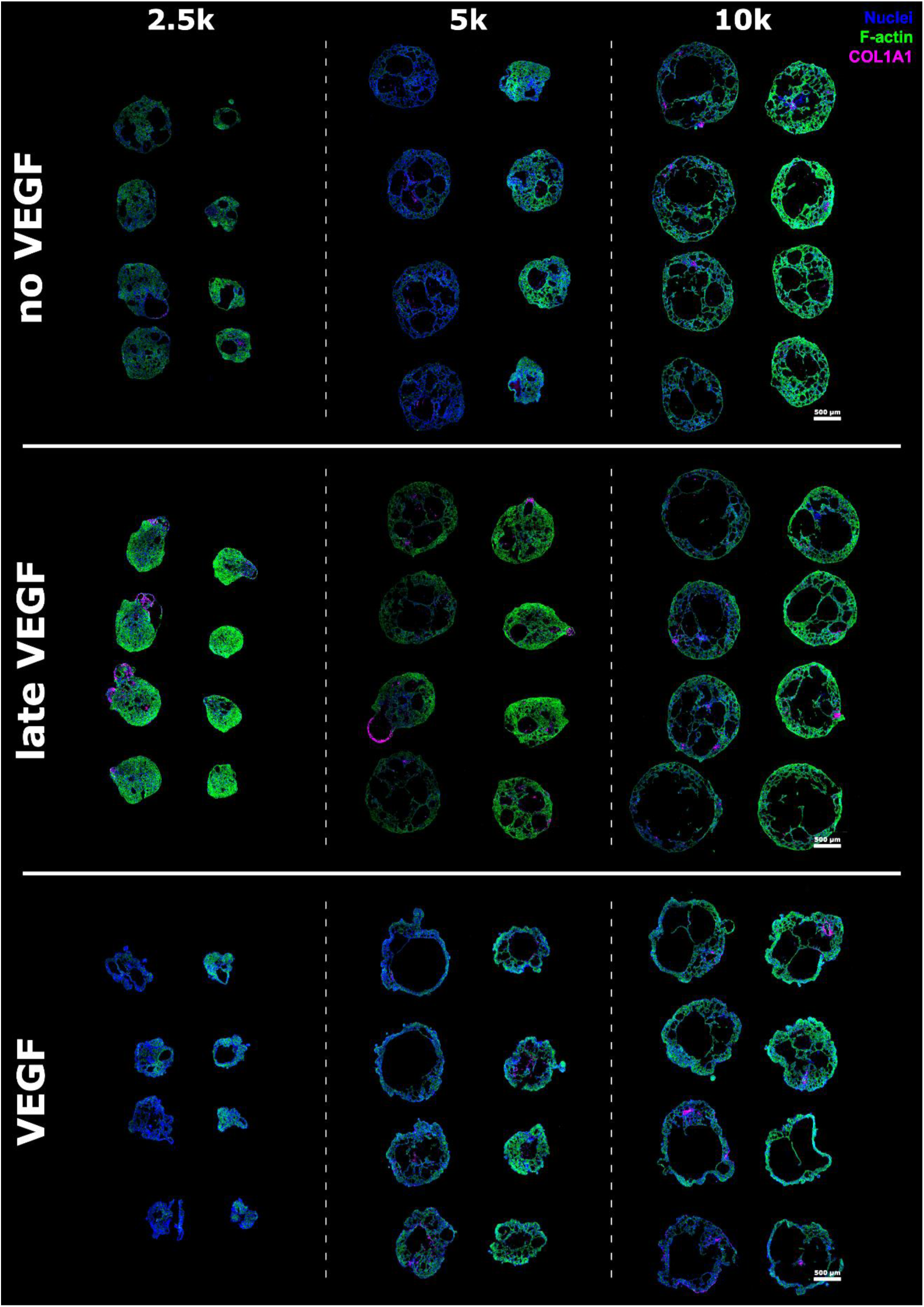
Immunohistochemical staining of cryosectioned cardioids (day 15) from each VEGF condition, generated using 2,500; 5,000 or 10,000 starting iPSCs. Nuclei (blue), F-actin (green), cTnT (magenta), COL1A1 (white). Scale bar: 500 µm.

**Figure S5.**
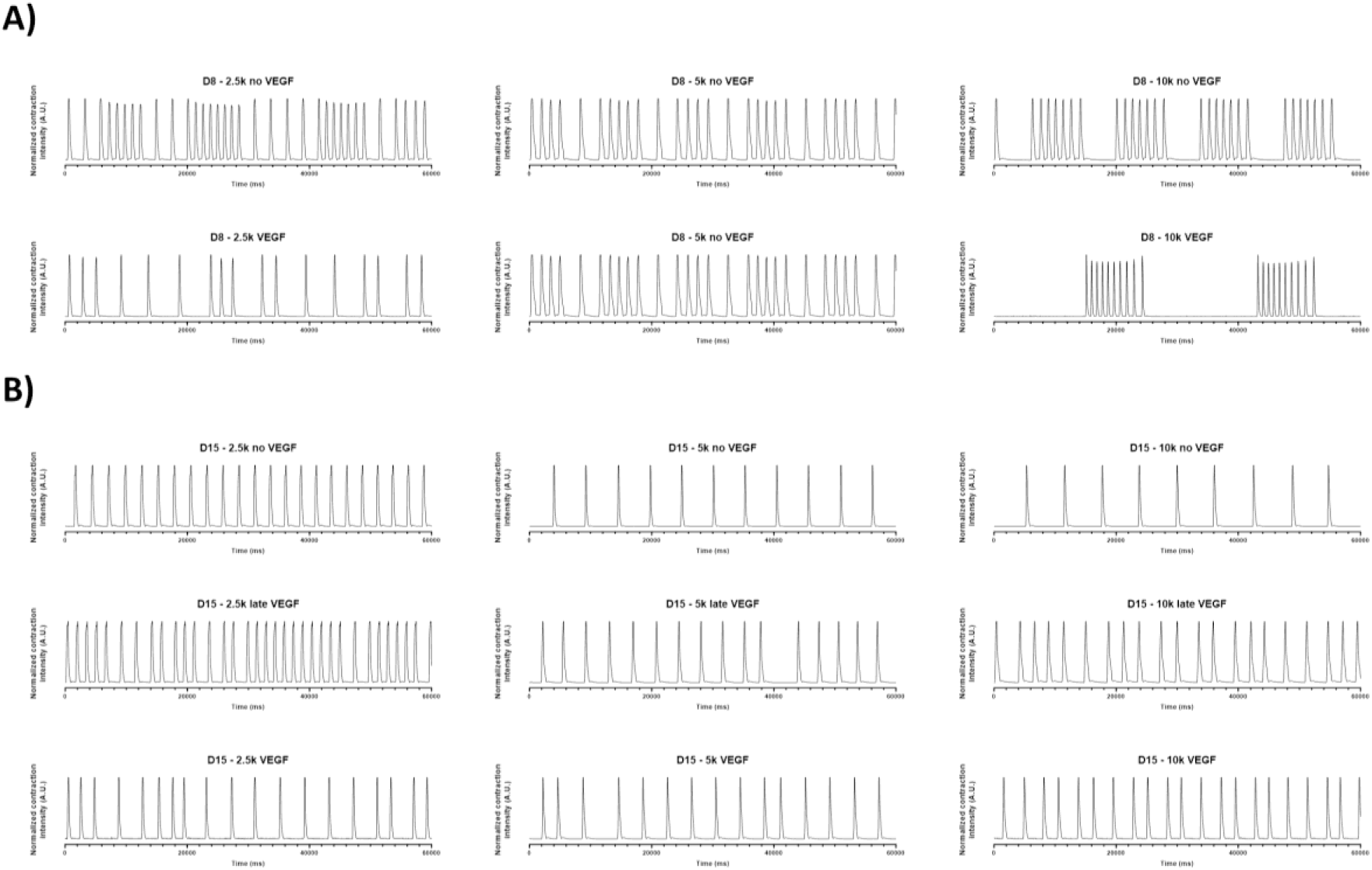
Normalized contraction profiles of non-dissociated cardioids formed using 2,500; 5,000 or 10,000 starting iPSCs and different VEGF conditions, at day 8 (A) and day 15 (B).

**Figure S6.**
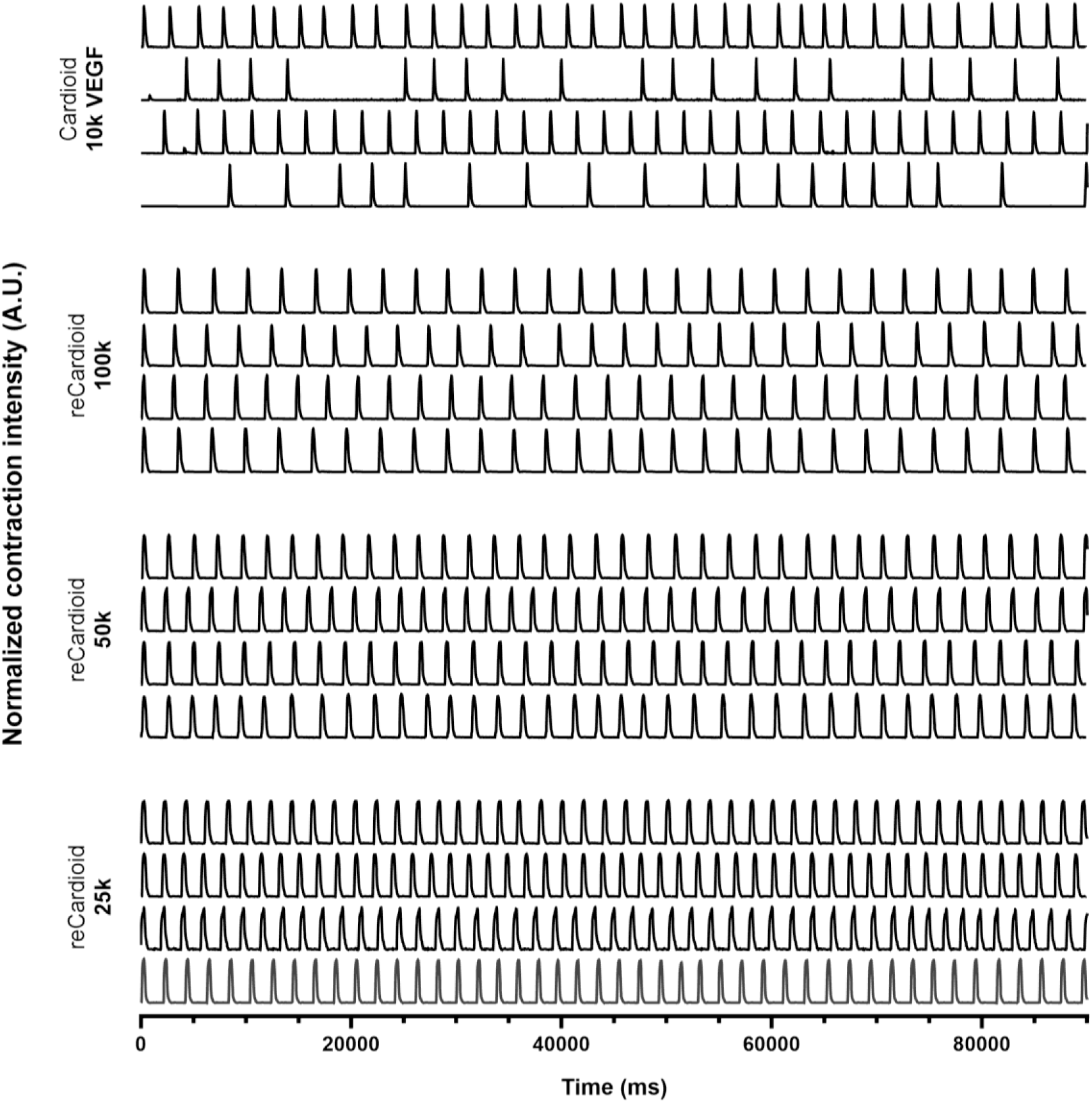
Normalized contraction profiles of original cardioids (VEGF condition and formed using 10,000 iPSCs) and reCardioids (formed using 25,000; 50,000 or 10,000 cells).

**Figure S7.**
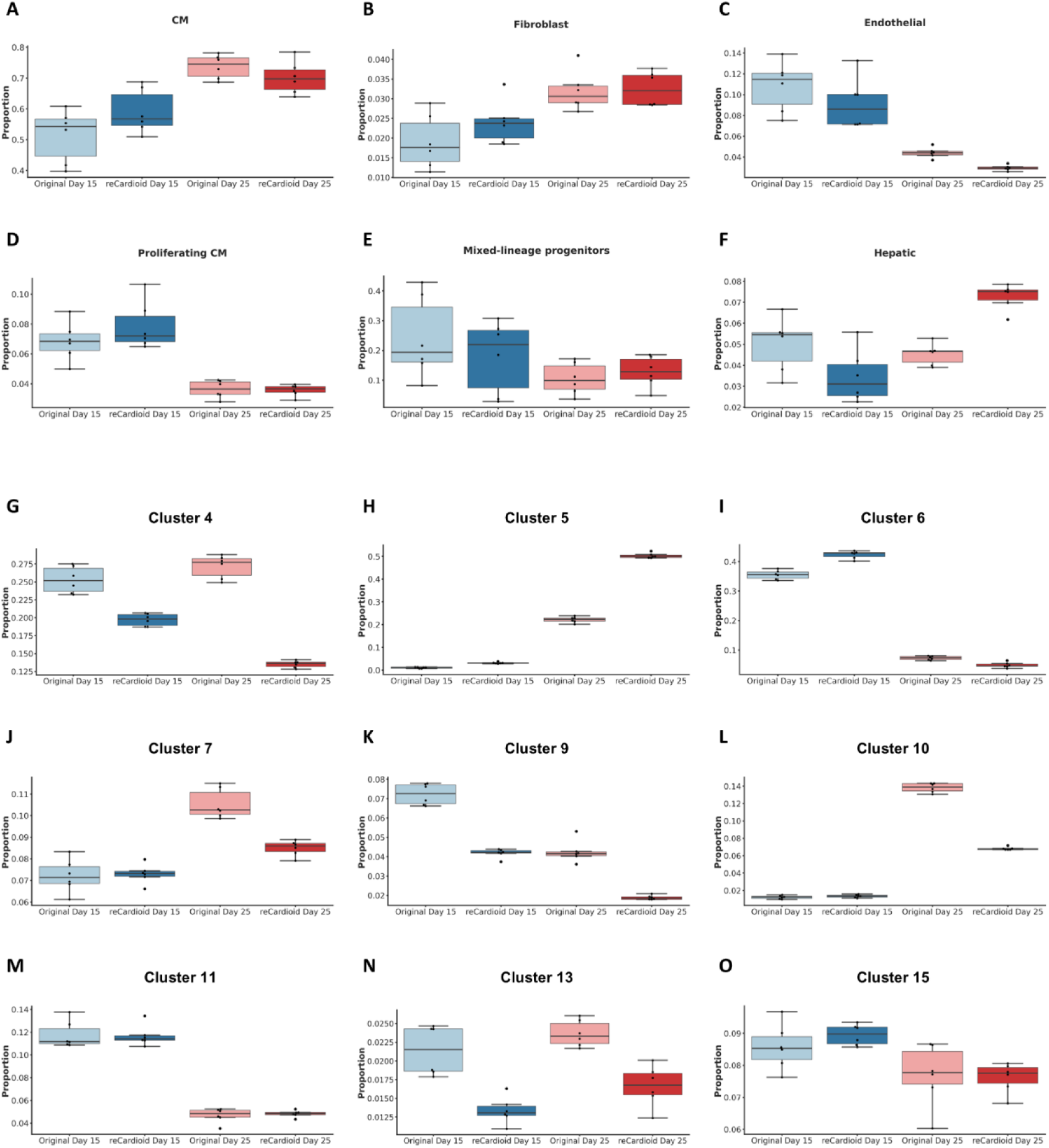
Quantification of cell type and subpopulation proportions across differentiation protocols and timepoints. (A-F) Boxplots displaying the proportion of identified major cell types: (A) CM (Cardiomyocytes), (B) Fibroblasts, (C) Endothelial cells, (D) Proliferating CMs, (E) Mixed-lineage progenitors, and (F) Hepatic cells. (G-O) Boxplots displaying the proportional breakdown of individual cardiomyocyte clusters: (G) Cluster 4, (H) Cluster 5, (I) Cluster 6, (J) Cluster 7, (K) Cluster 9, (L) Cluster 10, (M) Cluster 11, (N) Cluster 13, and (O) Cluster 15. All boxplots compare original non-dissociated cardioids and reCardioids at day 15 and day 25.

## References

[1] H. Sung, J. Ferlay, R. L. Siegel, M. Laversanne, I. Soerjomataram, A. Jemal, F. Bray, CA: A Cancer Journal for Clinicians 2021, 71, 209.

[2] K. D. Miller, L. Nogueira, T. Devasia, A. B. Mariotto, K. R. Yabroff, A. Jemal, J. Kramer, R. L. Siegel, CA: A Cancer Journal for Clinicians 2022, 72, 409.

[3] A. R. Lyon, T. López-Fernández, L. S. Couch, R. Asteggiano, M. C. Aznar, J. Bergler-Klein, G. Boriani, D. Cardinale, R. Cordoba, B. Cosyns, et al., Eur Heart J 2022, 43, 4229.

[4] J. J. Moslehi, New England Journal of Medicine 2016, 375, 1457.

[5] S. H. Armenian, C. J. Gibson, R. C. Rockne, K. K. Ness, J Natl Cancer Inst 2019, 111, 226.

[6] K. K. Ness, G. T. Armstrong, M. Kundu, C. L. Wilson, T. Tchkonia, J. L. Kirkland, Cancer 2015, 121, 1540.

[7] A. N. Linders, I. B. Dias, T. López Fernández, C. G. Tocchetti, N. Bomer, P. Van der Meer, npj Aging 2024, 10, 9.

[8] S. M. Swain, F. S. Whaley, M. S. Ewer, Cancer 2003, 97, 2869.

[9] J. V. McGowan, R. Chung, A. Maulik, I. Piotrowska, J. M. Walker, D. M. Yellon, Cardiovasc Drugs Ther 2017, 31, 63.

[10] S. C. Darby, M. Ewertz, P. McGale, A. M. Bennet, U. Blom-Goldman, D. Brønnum, C. Correa, D. Cutter, G. Gagliardi, B. Gigante, et al., New England Journal of Medicine 2013, 368, 987.

[11] J. D. Mitchell, D. A. Cehic, M. Morgia, C. Bergom, J. Toohey, P. A. Guerrero, M. Ferencik, R. Kikuchi, J. R. Carver, V. G. Zaha, et al., JACC: CardioOncology 2021, 3, 360.

[12] M. Kearney, M. Keys, C. Faivre-Finn, Z. Wang, M. C. Aznar, F. Duane, Radiotherapy and Oncology 2022, 172, 118.

[13] N. K. Taunk, B. G. Haffty, J. B. Kostis, S. Goyal, Front. Oncol. 2015, 5, DOI 10.3389/fonc.2015.00039.

[14] E. Belzile-Dugas, M. J. Eisenberg, Journal of the American Heart Association 2021, 10, e021686.

[15] K. C. Oeffinger, A. C. Mertens, C. A. Sklar, T. Kawashima, M. M. Hudson, A. T. Meadows, D. L. Friedman, N. Marina, W. Hobbie, N. S. Kadan-Lottick, et al., New England Journal of Medicine 2006, 355, 1572.

[16] S. H. Armenian, G. T. Armstrong, G. Aune, E. J. Chow, M. J. Ehrhardt, B. Ky, J. Moslehi, D. A. Mulrooney, P. C. Nathan, T. D. Ryan, et al., J Clin Oncol 2018, 36, 2135.

[17] M. Y. Desai, S. Windecker, P. Lancellotti, J. J. Bax, B. P. Griffin, O. Cahlon, D. R. Johnston, JACC 2019, 74, 905.

[18] E. Giacomelli, V. Meraviglia, G. Campostrini, A. Cochrane, X. Cao, R. W. J. van Helden, A. Krotenberg Garcia, M. Mircea, S. Kostidis, R. P. Davis, et al., Cell Stem Cell 2020, 26, 862.

[19] S. Tapio, J Radiat Res 2016, 57, 439.

[20] J. Wei, Q. Zhao, C. Lu, C. Gao, Z. Mu, D. Dong, M. Sun, Biochemical Pharmacology 2026, 247, 117745.

[21] A. Asnani, J. J. Moslehi, B. B. Adhikari, A. H. Baik, A. M. Beyer, R. A. de Boer, A. Ghigo, I. M. Grumbach, S. Jain, H. Zhu, et al., Circulation Research 2021, 129, e21.

[22] G. A. Van Norman, JACC: Basic to Translational Science 2019, 4, 845.

[23] V. G. Desai, E. H. Herman, C. L. Moland, W. S. Branham, S. M. Lewis, K. J. Davis, N. I. George, T. Lee, S. Kerr, J. C. Fuscoe, Toxicology and Applied Pharmacology 2013, 266, 109.

[24] E. Y. Podyacheva, E. A. Kushnareva, A. A. Karpov, Y. G. Toropova, Front. Pharmacol. 2021, 12, DOI 10.3389/fphar.2021.670479.

[25] Y. R. Lewis-Israeli, A. H. Wasserman, M. A. Gabalski, B. D. Volmert, Y. Ming, K. A. Ball, W. Yang, J. Zou, G. Ni, N. Pajares, et al., Nature Communications 2021, 12, 5142.

[26] P. Hofbauer, S. M. Jahnel, N. Papai, M. Giesshammer, A. Deyett, C. Schmidt, M. Penc, K. Tavernini, N. Grdseloff, C. Meledeth, et al., cell 2021, 184, 3299.

[27] C. Schmidt, A. Deyett, T. Ilmer, S. Haendeler, A. Torres Caballero, M. Novatchkova, M. A. Netzer, L. Ceci Ginistrelli, E. Mancheno Juncosa, T. Bhattacharya, et al., Cell 2023, 186, 5587.

[28] L. Drakhlis, S. Biswanath, C.-M. Farr, V. Lupanow, J. Teske, K. Ritzenhoff, A. Franke, F. Manstein, E. Bolesani, H. Kempf, et al., Nature Biotechnology 2021, 39, 737.

[29] A. C. Silva, O. B. Matthys, D. A. Joy, M. A. Kauss, V. Natarajan, M. H. Lai, D. Turaga, A. P. Blair, M. Alexanian, B. G. Bruneau, et al., Cell Stem Cell 2021, 28, 2137.

[30] A. B. Meier, D. Zawada, M. T. De Angelis, L. D. Martens, G. Santamaria, S. Zengerle, M. Nowak-Imialek, J. Kornherr, F. Zhang, Q. Tian, et al., Nat Biotechnol 2023, 41, 1787.

[31] D. J. Richards, Y. Li, C. M. Kerr, J. Yao, G. C. Beeson, R. C. Coyle, X. Chen, J. Jia, B. Damon, R. Wilson, et al., Nat Biomed Eng 2020, 4, 446.

[32] R. J. Mills, B. L. Parker, G. A. Quaife-Ryan, H. K. Voges, E. J. Needham, A. Bornot, M. Ding, H. Andersson, M. Polla, D. A. Elliott, et al., Cell Stem Cell 2019, 24, 895.

[33] R. J. Mills, D. M. Titmarsh, X. Koenig, B. L. Parker, J. G. Ryall, G. A. Quaife-Ryan, H. K. Voges, M. P. Hodson, C. Ferguson, L. Drowley, et al., Proc Natl Acad Sci U S A 2017, 114, E8372.

[34] X. Cao, D. Thomas, L. A. Whitcomb, M. Wang, A. Chatterjee, A. J. Chicco, M. M. Weil, J. C. Wu, Journal of Molecular and Cellular Cardiology 2024, 188, 105.

[35] T. Smit, E. Schickel, O. Azimzadeh, C. von Toerne, O. Rauh, S. Ritter, M. Durante, I. S. Schroeder, Cells 2021, 10, DOI 10.3390/cells10102608.

[36] D. N. Tavakol, T. R. Nash, Y. Kim, S. He, S. Fleischer, P. L. Graney, J. A. Brown, M. Liberman, M. Tamargo, A. Harken, et al., Biomaterials 2023, 301, 122267.

[37] H. Cao, L. Yue, J. Shao, F. Kong, S. Liu, H. Huai, Z. He, Z. Mao, Y. Yang, Y. Tan, et al., Stem Cell Res Ther 2024, 15, 493.

[38] F. Antonelli, International Journal of Molecular Sciences 2023, 24, DOI 10.3390/ijms241310620.

[39] J. Yi, L. Yue, Y. Zhang, N. Tao, H. Duan, L. Lv, Y. Tan, H. Wang, American Journal of Physiology-Cell Physiology 2023, 324, C1320.

[40] J. S. Kim, S. W. Choi, Y.-G. Park, S. J. Kim, C. H. Choi, M.-J. Cha, J. H. Chang, International Journal of Molecular Sciences 2021, 23, DOI 10.3390/ijms23010351.

[41] K. Livingston, R. A. Schlaak, L. L. Puckett, C. Bergom, Front. Cardiovasc. Med. 2020, 7, DOI 10.3389/fcvm.2020.00020.

[42] H. Kim, R. D. Kamm, G. Vunjak-Novakovic, J. C. Wu, Cell Stem Cell 2022, 29, 503.

[43] A. Arefin, M. Mendoza, K. Dame, M. I. Garcia, D. G. Strauss, A. J. S. Ribeiro, Front. Pharmacol. 2023, 14, DOI 10.3389/fphar.2023.1212092.

[44] T. de Korte, B. B. Johnson, G. Kosmidis, B. Samson-Couterie, M. P. H. Mol, R. W. J. van Helden, E. Razaghi, L. François, V. Meraviglia, L. Yiangou, et al., Trends in Biotechnology 2025, 0, DOI 10.1016/j.tibtech.2025.11.016.

[45] S.-J. Lee, H.-A. Lee, Organoid 2024, 4, e6.

[46] S. M. Biendarra-Tiegs, F. J. Secreto, T. J. Nelson, in Cell Biology and Translational Medicine, Volume 6: Stem Cells: Their Heterogeneity, Niche and Regenerative Potential (Ed: K. Turksen), Springer International Publishing, Cham, 2020, pp. 1–29.

[47] J. S. Reyat, Y. Shen, G. Poologasundarampillai, A. Moetazedian, J. Rayes, A. O. Khan, Circulation Research 2025, 137, 1133.

[48] S. Cho, D. E. Discher, K. W. Leong, G. Vunjak-Novakovic, J. C. Wu, Nat Methods 2022, 19, 1064.

[49] G. Rossi, N. Broguiere, M. Miyamoto, A. Boni, R. Guiet, M. Girgin, R. G. Kelly, C. Kwon, M. P. Lutolf, Cell Stem Cell 2021, 28, 230.

[50] P.-Y. Liang, Y. Chang, G. Jin, X. Lian, X. Bao, Front. Bioeng. Biotechnol. 2022, 10, DOI 10.3389/fbioe.2022.1059243.

[51] A. Guo, X. Zhang, V. R. Iyer, B. Chen, C. Zhang, W. J. Kutschke, R. M. Weiss, C. Franzini-Armstrong, L.-S. Song, Proceedings of the National Academy of Sciences 2014, 111, 12240.

[52] C. R. McKeown, R. B. Nowak, J. Moyer, M. A. Sussman, V. M. Fowler, Circulation Research 2008, 103, 1241.

[53] S. Yamashiro, D. S. Gokhin, S. Kimura, R. B. Nowak, V. M. Fowler, Cytoskeleton 2012, 69, 337.

[54] Y. Guo, W. T. Pu, Circulation Research 2020, 126, 1086.

[55] E. Karbassi, A. Fenix, S. Marchiano, N. Muraoka, K. Nakamura, X. Yang, C. E. Murry, Nat Rev Cardiol 2020, 17, 341.

[56] J. P. Goetze, B. G. Bruneau, H. R. Ramos, T. Ogawa, M. K. de Bold, A. J. de Bold, Nat Rev Cardiol 2020, 17, 698.

[57] L. B. Daniels, A. S. Maisel, JACC 2007, 50, 2357.

[58] W. Yan, F. Wu, J. Morser, Q. Wu, Proc. Natl. Acad. Sci. U.S.A. 2000, 97, 8525.

[59] N. Dong, S. Chen, J. Yang, L. He, P. Liu, D. Zheng, L. Li, Y. Zhou, C. Ruan, E. Plow, et al., Circulation: Heart Failure 2010, 3, 207.

[60] R. M. Vejandla, B.-O. Orgil, N. R. Alberson, N. Li, U. Munkhsaikhan, Z. Khuchua, R. Martherus, E. U. Azeloglu, F. Xu, L. Lu, et al., American Journal of Physiology-Heart and Circulatory Physiology 2021, 320, H2130.

[61] H. Wang, Z. Li, J. Wang, K. Sun, Q. Cui, L. Song, Y. Zou, X. Wang, X. Liu, R. Hui, et al., American Journal of Human Genetics 2010, 87, 687.

[62] C. Liu, S. Spinozzi, J.-Y. Chen, X. Fang, W. Feng, G. Perkins, P. Cattaneo, N. Guimarães-Camboa, N. D. Dalton, K. L. Peterson, et al., Circulation 2019, 140, 55.

[63] L. Kreuder, P.-A. Bissey, K. W. Yip, F.-F. Liu, International Journal of Radiation Biology 2025, 0, 1.

[64] J. Xuan, J. Zhou, Y. Huang, M. Huang, S. Pu, L. Fang, Y. Chen, J. Xue, Sci Rep 2026, DOI 10.1038/s41598-026-42367-5.

[65] K. Xue, J. Zhang, C. Li, J. Li, C. Wang, Q. Zhang, X. Chen, X. Yu, L. Sun, X. Yu, Journal of Cellular and Molecular Medicine 2019, 23, 4229.

[66] B. Baselet, P. Sonveaux, S. Baatout, A. Aerts, Cell. Mol. Life Sci. 2019, 76, 699.

[67] C. E. Friedman, Q. Nguyen, S. W. Lukowski, A. Helfer, H. S. Chiu, J. Miklas, S. Levy, S. Suo, J.-D. J. Han, P. Osteil, et al., Cell Stem Cell 2018, 23, 586.

[68] F. Capone, A. Vacca, G. Bidault, D. Sarver, D. Kaminska, S. Strocchi, A. Vidal-Puig, C. M. Greco, A. J. Lusis, G. G. Schiattarella, Circulation Research 2025, 136, 1335.

[69] A. Hakeem, J. van Berlo, X. S. Revelo, JACC: Basic to Translational Science 2025, 10, 101309.

[70] K. Ronaldson-Bouchard, D. Teles, K. Yeager, D. N. Tavakol, Y. Zhao, A. Chramiec, S. Tagore, M. Summers, S. Stylianos, M. Tamargo, et al., Nat Biomed Eng 2022, 6, 351.

[71] T. Magdy, A. J. T. Schuldt, J. C. Wu, D. Bernstein, P. W. Burridge, Annual Review of Pharmacology and Toxicology 2018, 58, 83.

[72] D. A. M. Feyen, W. L. McKeithan, A. A. N. Bruyneel, S. Spiering, L. Hörmann, B. Ulmer, H. Zhang, F. Briganti, M. Schweizer, B. Hegyi, et al., Cell Reports 2020, 32, 107925.

[73] Y. Guo, B. D. Jardin, P. Zhou, I. Sethi, B. N. Akerberg, C. N. Toepfer, Y. Ai, Y. Li, Q. Ma, S. Guatimosim, et al., Nat Commun 2018, 9, 3837.

[74] S. S. Nunes, J. W. Miklas, J. Liu, R. Aschar-Sobbi, Y. Xiao, B. Zhang, J. Jiang, S. Masse, M. Gagliardi, A. Hsieh, et al., Nat Methods 2013, 10, 781.

[75] J.-L. Ruan, N. L. Tulloch, M. V. Razumova, M. Saiget, V. Muskheli, L. Pabon, H. Reinecke, M. Regnier, C. E. Murry, Circulation 2016, 134, 1557.

[76] H. Tang, R. Liu, Z. Xiong, Q. He, G. Lu, W.-Y. Chan, W. Wang, 2025.

[77] H. Sawaya, I. A. Sebag, J. C. Plana, J. L. Januzzi, B. Ky, V. Cohen, S. Gosavi, J. R. Carver, S. E. Wiegers, R. P. Martin, et al., The American Journal of Cardiology 2011, 107, 1375.

[78] S. Martel, C. Maurer, M. Lambertini, N. Pondé, E. De Azambuja, Expert Opinion on Drug Safety 2017, 16, 1021.

[79] E. T. H. Yeh, C. L. Bickford, Journal of the American College of Cardiology 2009, 53, 2231.

[80] X. Han, Y. Zhou, W. Liu, npj Precision Onc 2017, 1, 31.

[81] J. Alexandre, J. Cautela, S. Ederhy, G. L. Damaj, J. Salem, F. Barlesi, L. Farnault, A. Charbonnier, M. Mirabel, S. Champiat, et al., Journal of the American Heart Association 2020, 9, e018403.

[82] P. S. Rawat, A. Jaiswal, A. Khurana, J. S. Bhatti, U. Navik, Biomedicine & Pharmacotherapy 2021, 139, 111708.

[83] C. G. Nebigil, L. Désaubry, Front. Pharmacol. 2018, 9, DOI 10.3389/fphar.2018.01262.

[84] A. Boyd, P. Stoodley, D. Richards, R. Hui, P. Harnett, K. Vo, T. Marwick, L. Thomas, PLOS ONE 2017, 12, e0175544.

[85] M. Arzt, B. Gao, M. Mozneb, S. Pohlman, R. B. Cejas, Q. Liu, F. Huang, C. Yu, Y. Zhang, X. Fan, et al., Stem Cell Reports 2023, 18, 1913.

[86] C. Liu, M. Shen, Y. Liu, A. Manhas, S. R. Zhao, M. Zhang, N. Belbachir, L. Ren, J. Z. Zhang, A. Caudal, et al., Cell Stem Cell 2024, 31, 1760.

[87] X. Chen, N. Lu, S. Huang, Y. Zhang, Z. Liu, X. Wang, Chemico-Biological Interactions 2023, 386, 110777.

[88] J. Jang, H. Jung, J. Jeong, J. Jeon, K. Lee, H. R. Jang, J.-W. Han, J. Lee, Heliyon 2024, 10, e38714.

[89] S. P. Jackson, J. Bartek, Nature 2009, 461, 1071.

[90] A. V. Gudkov, E. A. Komarova, Nat Rev Cancer 2003, 3, 117.

[91] S. Xie, S.-C. Xu, W. Deng, Q. Tang, Sig Transduct Target Ther 2023, 8, 114.

[92] I. J. Goldberg, C. M. Trent, P. C. Schulze, Cell Metabolism 2012, 15, 805.

[93] T. Doenst, T. D. Nguyen, E. D. Abel, Circulation Research 2013, 113, 709.

[94] G. D. Lopaschuk, Q. G. Karwi, R. Tian, A. R. Wende, E. D. Abel, Circulation Research 2021, 128, 1487.

[95] B. C. Bernardo, K. L. Weeks, L. Pretorius, J. R. McMullen, Pharmacology & Therapeutics 2010, 128, 191.

[96] N. Frey, E. N. Olson, Annual Review of Physiology 2003, 65, 45.

[97] C. R. Dufour, B. J. Wilson, J. M. Huss, D. P. Kelly, W. A. Alaynick, M. Downes, R. M. Evans, M. Blanchette, V. Giguère, Cell Metabolism 2007, 5, 345.

[98] X. Wu, P. Eder, B. Chang, J. D. Molkentin, Proceedings of the National Academy of Sciences 2010, 107, 7000.

[99] J.-P. Coppé, P.-Y. Desprez, A. Krtolica, J. Campisi, Annual Review of Pathology: Mechanisms of Disease 2010, 5, 99.

[100] P. Ell, J. M. Martin, D. A. Cehic, D. T. M. Ngo, A. L. Sverdlov, Curr. Treat. Options in Oncol. 2021, 22, 70.

[101] M. D’Antonio, P. Benaglio, D. Jakubosky, W. W. Greenwald, H. Matsui, M. K. R. Donovan, H. Li, E. N. Smith, A. D’Antonio-Chronowska, K. A. Frazer, Cell Reports 2018, 24, 883.

[102] A. D. Panopoulos, M. D’Antonio, P. Benaglio, R. Williams, S. I. Hashem, B. M. Schuldt, C. DeBoever, A. D. Arias, M. Garcia, B. C. Nelson, et al., Stem Cell Reports 2017, 8, 1086.

[103] L. Sala, B. J. van Meer, L. G. J. Tertoolen, J. Bakkers, M. Bellin, R. P. Davis, C. Denning, M. A. E. Dieben, T. Eschenhagen, E. Giacomelli, et al., Circulation Research 2018, 122, e5.

